# Polyploidy is widespread in Microsporidia

**DOI:** 10.1101/2023.09.29.560119

**Authors:** Amjad Khalaf, Kamil S. Jaron, Mark L. Blaxter, Mara K. N. Lawniczak

## Abstract

Microsporidia are obligate intracellular eukaryotic parasites with an extremely broad host range. They have both economic and public health importance. Ploidy in microsporidia is variable, with a few species formally identified as diploid and one as polyploid. Given the increase in the number of studies sequencing microsporidian genomes, it is now possible to assess ploidy levels across all currently-explored microsporidian diversity. We estimate ploidy for all microsporidian datasets available on the Sequence Read Archive (SRA) using k-mer-based analyses, and demonstrate that polyploidy is widespread in Microsporidia, and that ploidy change is dynamic in the group. Using genome-wide heterozygosity estimates, we also show that polyploid microsporidian genomes are relatively homozygous, and we discuss the implications of these findings on the timing of polyploidisation events and their origin.

**Author Summary:** Microsporidia are single-celled intracellular parasites, distantly related to fungi, that can infect a broad range of hosts, from humans all the way to protozoans. Exploiting the wealth of microsporidian genomic data available, we use k-mer based analyses to assess ploidy status across the group. Understanding a genome’s ploidy is crucial in order to assemble it effectively, and may also be relevant for better understanding a parasite’s behaviour and life cycle. We show that tetraploidy is present in at least six species in Microsporidia, and that these polyploidisation events are likely to have occurred independently. We discuss why these findings may be paradoxical, given that Microsporidia, like other intracellular parasites, have extremely small, reduced genomes.

## Introduction

Microsporidia are eukaryotic, obligately intracellular, spore-forming parasites [1], with a broad range of hosts including protozoans, arthropods and vertebrates [2–5]. They were first discovered in 1857 as the cause of a disease known as “pébrine” in silkworms (*Bombyx mori*, Lepidoptera) which had decimated the silk industry from France to China [6–8]. In 2001, the first microsporidian genome was published, belonging to *Encephalitozoon cuniculi*, a parasite of the European rabbit (*Oryctolagus cuniculus*) used as a model in infection studies [9]. More recently, many studies have targeted microsporidian genomes, and microsporidia have become a valuable model for studying the evolution of parasitism [10–15]. Microsporidia have the smallest eukaryotic genomes, having undergone extreme reduction, losing many metabolic pathways, such as glycolysis and lipid biosynthesis [11,16–21] and lacking mitochondrial genomes because of reductive evolution of mitochondria to form mitosomes [22].

Understanding genome ploidy is critical for correct genome assembly and interpretation, and in Microsporidia is also central to understanding their life cycles and parasitism. However, microsporidian ploidy remains a major source of uncertainty. In part, this is due to the fact that microsporidians are thought to undergo cryptomitosis, where the condensation and separation of chromatin into distinct chromosomes is unclear [23,24]. This is further complicated by microsporidia being extremely small and difficult to culture, making isolation of their nuclei from their hosts challenging.

Given that Microsporidia are thought to descend from a sexual fungal lineage (related to the ancestor of Rozellomycota) [25], one possibility is that they retain, in general, ploidy similar to that of their ancestors. In fungi, sexual individuals cycle between different ploidy levels (haploid and diploid, or diploid and tetraploid). Two compatible partners with the lower ploidy level will recognise one another and fuse to form a zygote with the higher ploidy level. When triggered by the right conditions, the zygote undergoes meiosis to produce spores, which may be uni- or di-karyotic [26]. Like Basidomycota, some Microsporidia, including *Pleistophora debaisieuxi, Astathelohania contejeani, Ameson michaelis*, and *Toguebayea baccigeri* [27–31] also cycle between unikaryons and dikaryons (referred to as diplokaryons in Microsporidia), although we do not know if this is associated with ploidy change. Imaging of fixed material has shown evidence for the occurrence of putative events that mirror the fungal sexual cycle in Microsporidia such as gametogenesis, plasmogamy, karyogamy and meiosis [28,32–44].

In contrast, many microsporidian species have been reported to be strictly monokaryotic (e.g. *Encephalitozoon cuniculi*) or strictly diplokaryotic (e.g. *Vairimorpha ceranae* [previously known as *Nosema ceranae*]) for their full life cycle [28,32,45,46]. In three strictly monokaryotic species (*Encephalitozoon cuniculi, Nematocida parisii*, and *Nematocida sp1*) some heterozygosity was detected in a few genes across a small number of isolated strains, suggesting these species are diploid [9,47–50]. On the other hand, tetraploidy was recently shown to occur in a strictly diplokaryotic species (*Vairimorpha ceranae*) using k-mer based genome analysis [51]. The latter observation suggests that the microsporidian diplokaryon may behave like the morphologically similar diplomonad diplokaryon, as seen in *Giardia lamblia*, which cycles between tetraploid and octoploid states [52].

Nevertheless, all current observations relating to ploidy of Microsporidia are restricted to a few species, and collected using different methods. Given the wealth of microsporidian sequencing data now available, it is possible to assess ploidy levels across a wider diversity of taxa. Here we survey all microsporidian genomic sequencing datasets available publicly in the Sequence Read Archive (SRA) [53] using k-mer-based analyses to make a statement about ploidy across microsporidian phylogeny. We demonstrate that polyploidy is widespread in Microsporidia. By measuring heterozygosity within species at a genome-wide level, we show that the homeologous genomes within polyploids are relatively homozygous. We discuss whether polyploidy is a recent occurrence, and what processes may have given rise to it multiple times.

## Materials and Methods

### Download of NCBI SRA datasets

On the 7th of April 2023, we downloaded all microsporidian whole-genome DNA datasets available in the NCBI Sequence Read Archive (SRA) [53] using the SRA ToolKit ([version 3.0.3, SRA Toolkit Development Team 2023](https://github.com/ncbi/sra-tools)). This retrieved 217 samples from 46 different species (supplementary information Tables S1 and S2).

### Ploidy estimation

To estimate ploidy, we generated a k-mer spectrum for each sample using Jellyfish ([version 2.2.10](https://github.com/gmarcais/Jellyfish)) with k-mers of length 21 [54]. We analysed these k-mer spectra with Genomescope2 ([version 2.0](https://github.com/tbenavi1/genomescope2.0)) and Smudgeplot ([version 0.2.5](https://github.com/KamilSJaron/smudgeplot)) [55] for all the samples. Genomescope2 uses a k-mer spectrum from unassembled read sets to infer genome size and heterozygosity, and estimates ploidy. Smudgeplot is a visualisation technique that receives as input k-mer pairs where each k-mer is paired with those differing by one SNP. The k-mers in each pair are named A and B, with A being the k-mer with coverage greater than or equal to B. In a polyploid genome, k-mers A and B can be found in one or more genomic copies. For example, in a triploid, only the combination AAB can be found, but in a tetraploid, both AAAB and AABB are possible. Ploidy can be inferred by calculating the proportional coverage of B in relation to the total coverage in each k-mer structure combination (see [55]). To ensure that the ploidy estimates were reliable, we only considered datasets where the monoploid (1n) coverage was estimated to be over 20 fold, and a good model fit in GenomeScope2 was achieved. For a full explanation of how to interpret GenomeScope2 and Smudgeplot outputs, refer to the [Smudgeplot Github](https://github.com/KamilSJaron/smudgeplot) page or [55].

### Ribosomal small subunit (SSU) sequence reconstruction

For a representative readset of each of the 16 species surveyed (supplementary information Table S3), we used PhyloFlash ([version 3.3b1](http://hrgv.github.io/phyloFlash/), with the -emirge flag) to reconstruct an approximation of its ribosomal small subunit RNA (SSU) sequences [56,57]. We note that this process will collapse homeologous haplotypes and within-species variation. The identity of the reconstructed SSUs was confirmed by using NCBI Blast against NCBI NT [58].

### Phylogeny

We used the reconstructed SSU sequences for the 16 species for which we estimated ploidy, and an additional 116 SSU sequences from related microsporidian species on NCBI NT (supplementary information Table S4), to generate a phylogeny with *Rozella allomycis* (Rozellomycota) as the outgroup. The sequences were aligned using MAFFT (mafft -- maxiterate 1000 --globalpair) [59]. The phylogeny was inferred by generating a consensus maximum-likelihood tree using IQ-TREE (version 2.2.2.3) [60], with a GTR+F+I+R6 nucleotide substitution model, 1000 ultrafast bootstrap replicates, and 1000 bootstrap replicates for the SH-like approximate likelihood ratio test (iqtree -b 1000 -alrt 1000 -m GTR+F+I+R6). The model was chosen using IQ-TREE’s best-fit model finder according to Bayesian information criterion. The resultant tree was annotated using [Toytree](https://github.com/eaton-lab/toytree) and InkScape (version 1.2.2) to show the position of tetraploid and diploid species.

We also used the reconstructed SSU sequences for the 16 species for which we estimated ploidy, to generate a smaller phylogeny with *Rozella allomycis* (Rozellomycota) as the outgroup. We aligned the sequences using MAFFT (mafft --maxiterate 1000 --globalpair) [59]. The phylogeny was inferred by generating a consensus maximum-likelihood tree using IQ-TREE (version 2.2.2.3) [60], with the GTR+F+I+R6 nucleotide substitution model, 1000 ultrafast bootstrap replicates, and 1000 bootstrap replicates for the SH-like approximate likelihood ratio test as above. The alignment of this smaller dataset had better fit to a more parameterised model according to IQ-TREE’s best-fit model finder. However, we used the same model used in the larger alignment to maintain consistency. The resultant tree was annotated using [Toytree](https://github.com/eaton-lab/toytree) and InkScape (version 1.2.2) to show the position of tetraploid and diploid species.

## Results

### Polyploidy is widespread in Microsporidia

We estimated ploidy across 217 unassembled readsets from 46 species from the NCBI Sequence Read Archive (SRA) using Genomescope2 and Smudgeplot [55]. We filtered out samples with contamination, low coverage samples, and samples that consisted of dominant host peaks (Fig 1). We also filtered out three samples with unresolved Smudgeplot patterns that we were not able to interpret (Fig 2).

**Fig 1:**
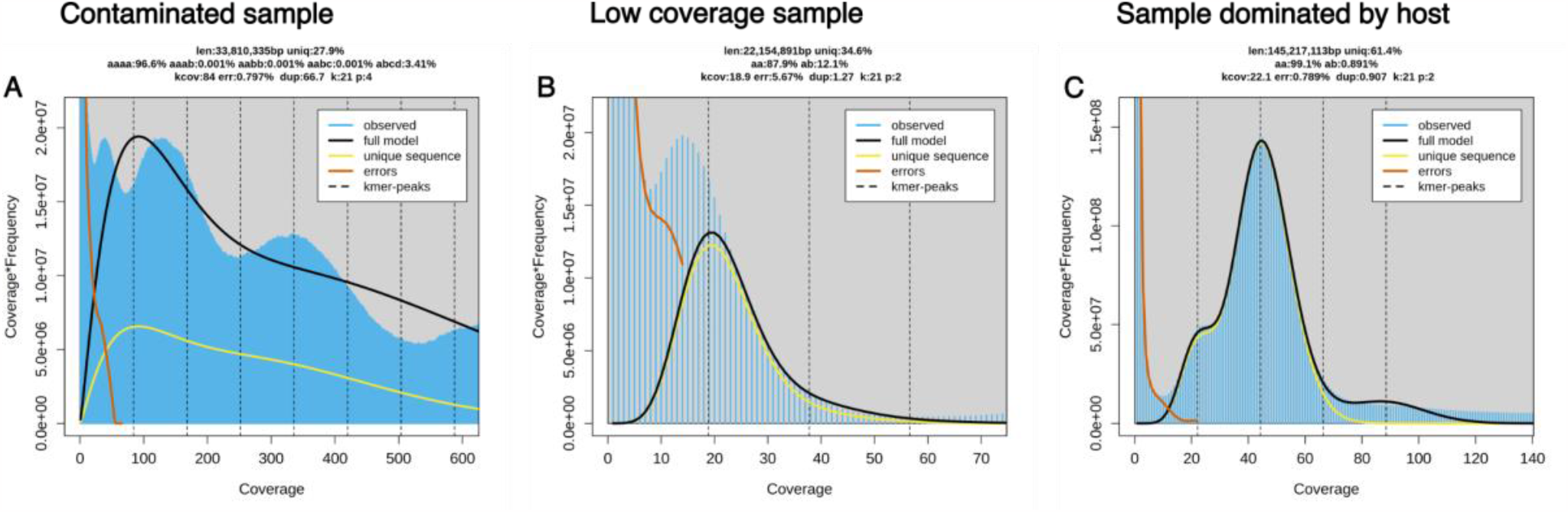
Examples of SRA datasets that were not analysed further. (A) GenomeScope2 transformed-linear plot of *Vairimorpha ceranae* (SRR7178080) as an example of a contaminated sample where the pattern coverage peaks are distorted by host contamination and a good model fit is not achieved. (B) GenomeScope2 transformed-linear plot of *Encephalitozoon cuniculi* (SRR122315) as an example of a low coverage sample. (C) GenomeScope2 transformed-linear plot of *Ordospora colligata* (SRR18286429) as an example of a sample that is dominated by host data, showing the predicted genome size to be ∼150 Mb.

**Fig 2:**
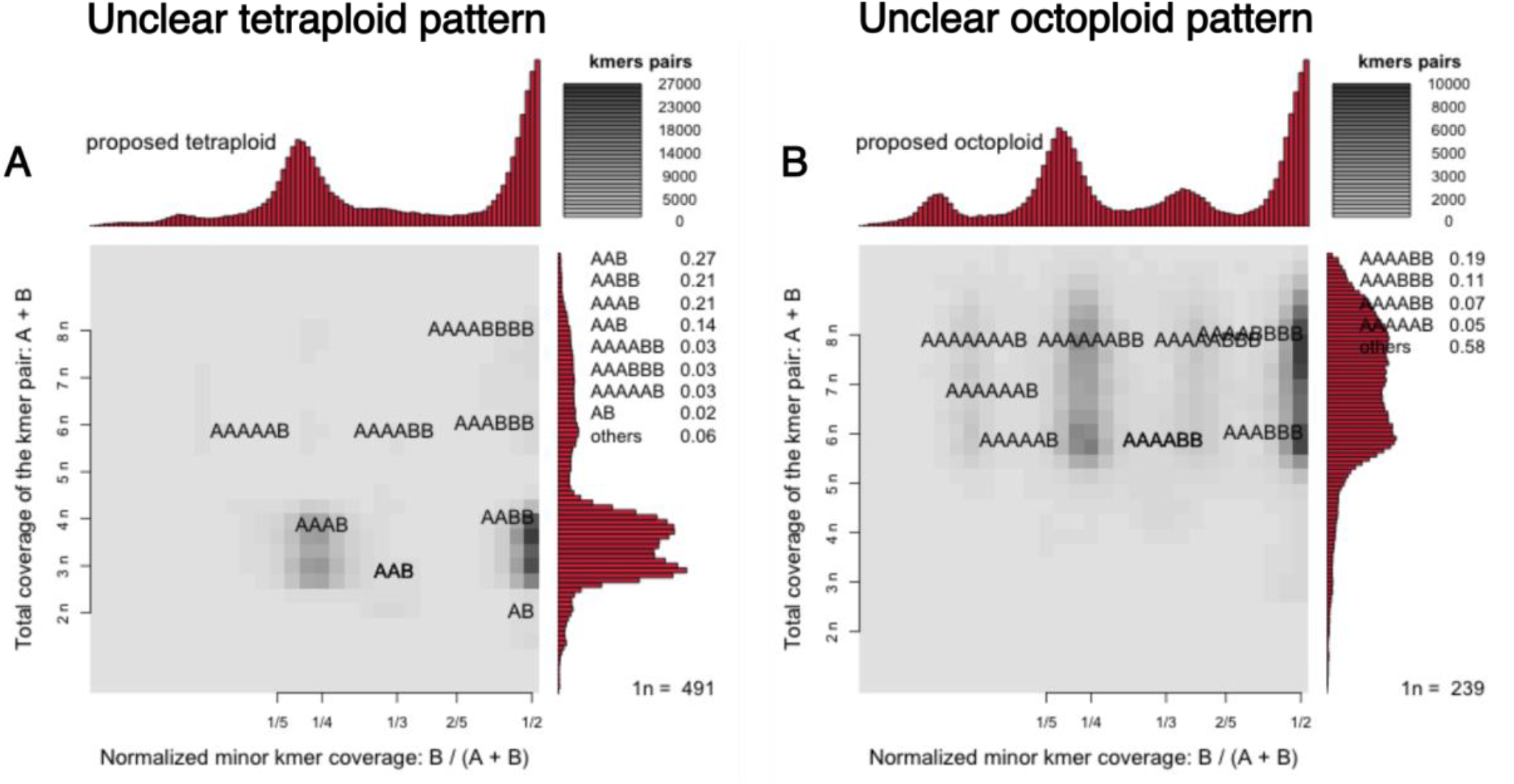
Samples with Smudgeplot patterns that were difficult to interpret. (A) *Vairimorpha ceranae* (SRR18590836) Smudgeplot with a smeared tetraploid pattern between 4n and 3n coverage. (B) *Vairimorpha ceranae* (SRR18590837) Smudgeplot with a smeared octoploid pattern between 6n and 8n coverage. These patterns remain difficult to interpret using current data.

After filtering, we retained 66 samples (from 16 species), all of which showed either diploid or tetraploid Smudgeplot patterns and varying levels of heterozygosity. For example, the GenomeScope2 plot for *Astathelohania contejeani* shows four peaks at 1n, 2n, 3n, and 4n coverage (Fig 3A), whilst the Smudgeplot shows strong signal at ¼ and ½ normalised minor k-mer coverage (Fig 3B). These patterns are indicative of tetraploidy. In contrast, the GenomeScope2 plot for *Nematocida ausubeli* shows only two peaks at 1n and 2n coverage (Fig 3C), whilst the Smudgeplot shows strong signal only at ½ normalised k-mer coverage (Fig 3D). This is indicative of diploidy. For the Encephalitozoonidae surveyed in this study, GenomeScope2 plots were interpreted as indicating a diploid genome with extremely reduced heterozygosity (<0.5%). For example, *Encephalitozoon intestinalis* (Fig 3E) had a a single, major k-mer coverage peak (∼400 fold in Fig 3E), and very low signals at half coverage (∼200 fold in Fig 3E) and double coverage (∼800 fold in Fig 3E). The Smudgeplot (Fig 3F) confirms highly homozygous diploidy, as some signal is seen at ½ normalised minor k-mer coverage with ∼400 fold (2n) coverage. The signal at ½ normalised minor k-mer coverage with ∼800 fold (4n) coverage is likely to derive from imperfect duplications in the genome. This pattern is not consistent with haploidy, triploidy, tetraploidy, or any other higher ploidies. Figures for all 66 samples are available in supporting information Table S1. Multiple samples, an average of four per species, were analysed for many species (supplementary information Table S1). Where multiple samples were available for a nominal species, they showed a consistent ploidy signal.

**Fig 3:**
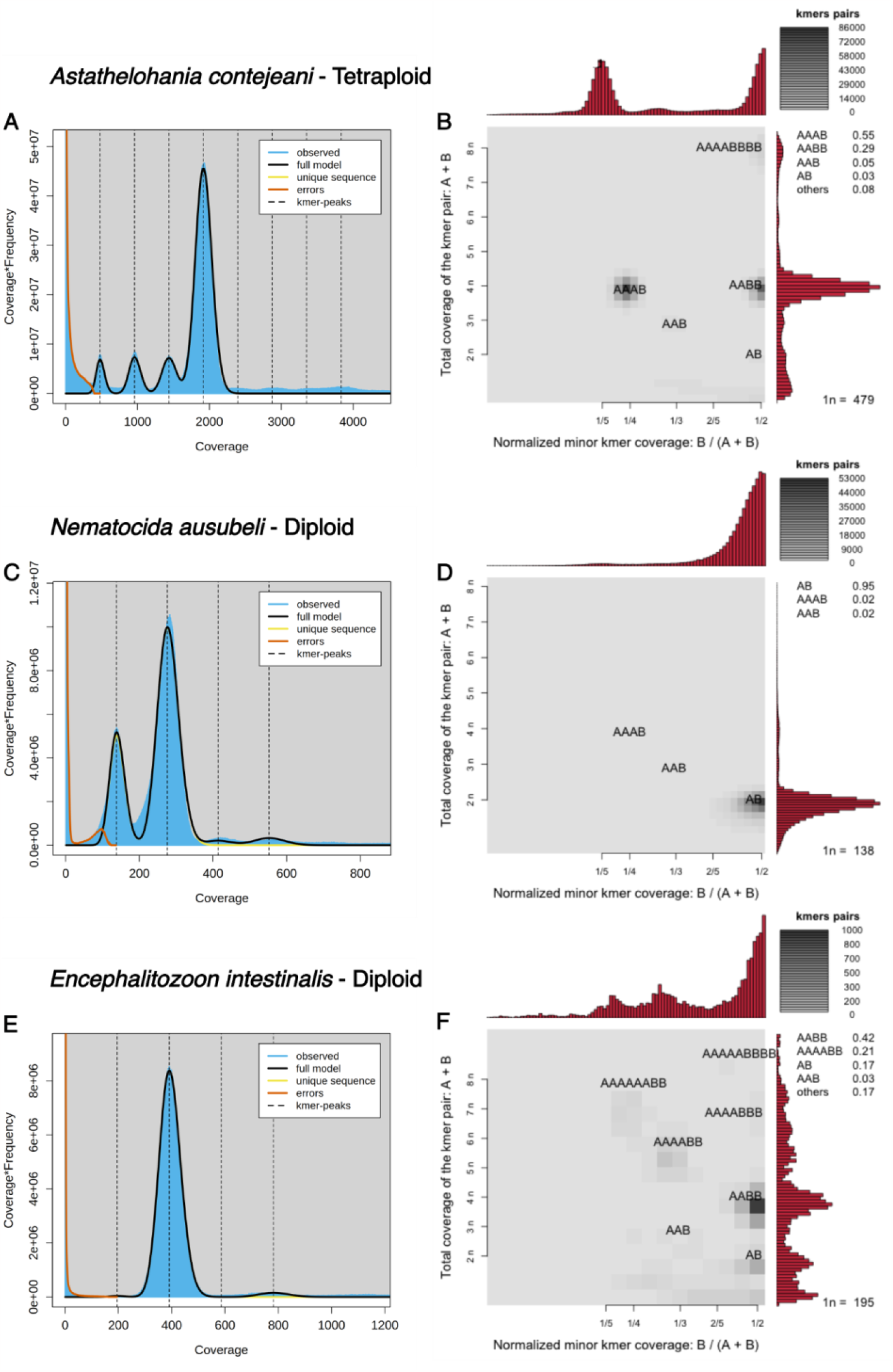
Ploidy estimates for Microsporidia species. (A) and (B) GenomeScope2 and Smudgeplot results for *Astathelohania contejeani* (SRR8476226) estimating tetraploidy. (C) and (D) GenomeScope2 and Smudgeplot results for *Nematocida ausubeli* (SRR350188) estimating diploidy. (E) and (F) GenomeScope2 and Smudgeplot results for *Encephalitozoon intestinalis* (SRR24007516) estimating diploidy with exceptionally high homozygosity.

Mapping the ploidy estimates to the phylogeny estimated from the SSU locus demonstrated that only diploid species were identified in the clade Ovavesiculida, and no data were available for the clade Amblyosporida. We identified six tetraploid species from three of the six major microsporidian clades (Nosematida, Glugeida, Neopereziida), and the orphan lineage represented by *Astathelohania contejeani* [12]. Neither diploidy nor tetraploidy was monophyletic on the phylogeny. This implies multiple events of gain of polyploidy (five events, if the last common ancestor of Microsporidia is assumed to have been diploid, and only tetraploidy gains are permitted) or gain and loss of polyploidy (for example one gain and three losses if the last common ancestor of the clade defined by *Astathelohania* and *Varimorpha* gained tetraploidy) (Figs 4 and 5).

**Fig 4:**
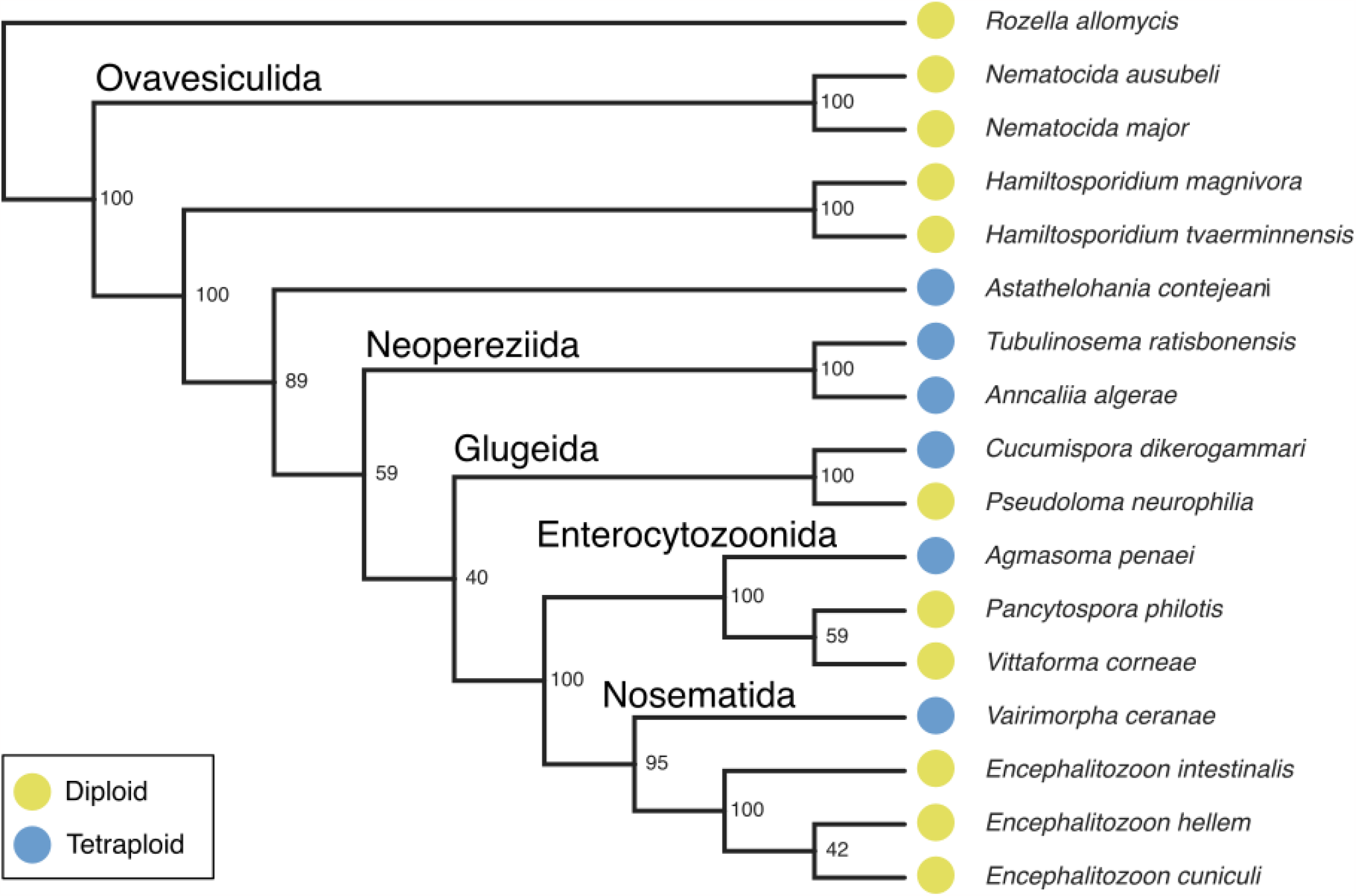
A cladogram of microsporidian species with confident ploidy estimates, rooted on *Rozella allomycis* (Diploid, [61]). The cladogram was inferred from ribosomal small subunit sequences using IQ-TREE (version 2.2.2.3), with a GTR+F+I+R6 nucleotide substitution model.

**Fig 5:**
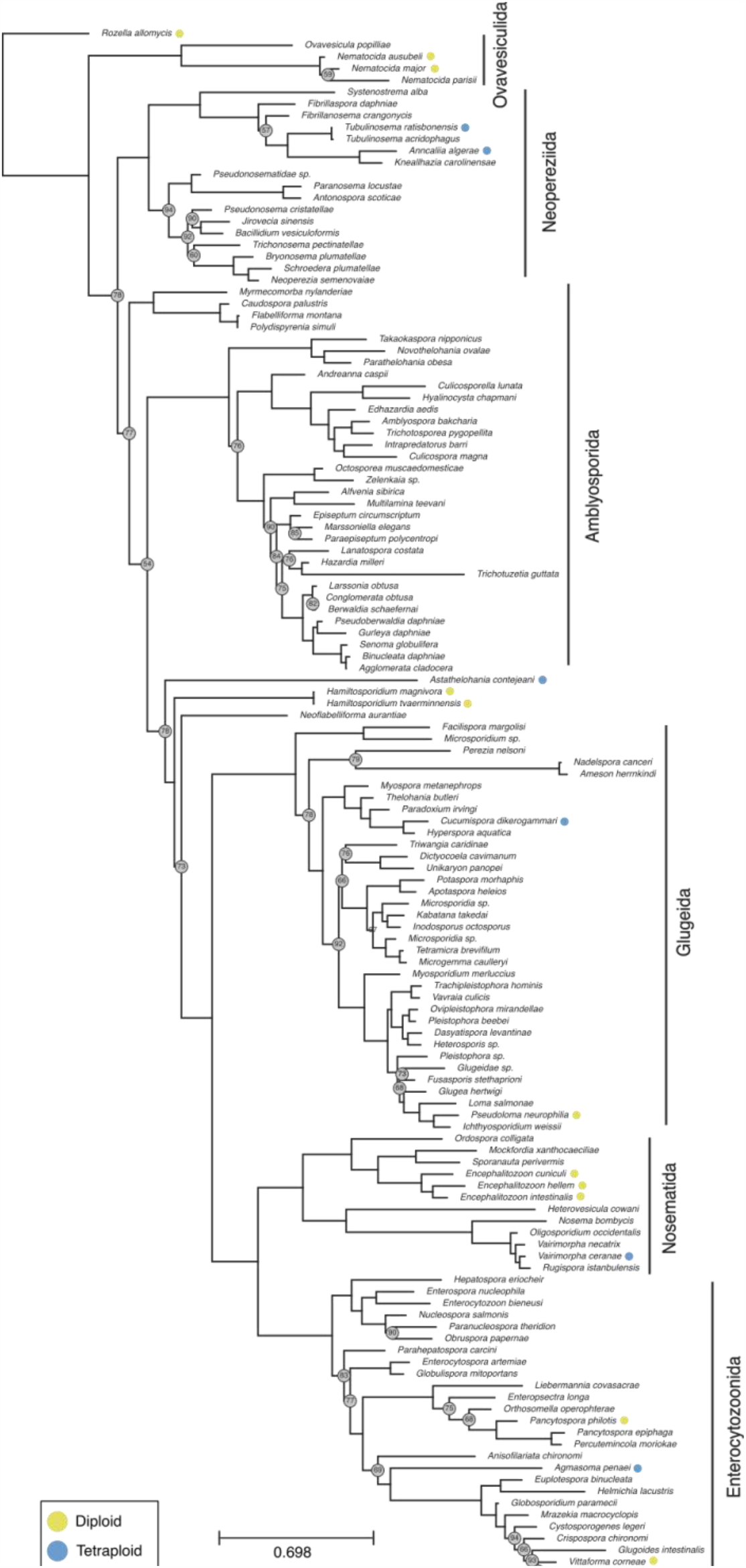
Phylogeny of Microsporidia, rooted on *Rozella allomycis* (Diploid, [61]). The phylogeny was inferred from ribosomal small subunit sequences using IQ-TREE (version 2.2.2.3), with a GTR+F+I+R6 nucleotide substitution model. Nodes with bootstrap values<95% are indicated with grey circles.

### Most tetraploid genomes are more than 96% homozygous

We estimated genome-wide heterozygosity across all samples of the 16 species for which we were able to estimate ploidy. Genome-wide heterozygosity in tetraploids varied almost eight-fold, ranging from an average of ∼8% in *Agmasoma penaei* to an average of ∼1% in *Vairimorpha ceranae*. In each sample, we also estimated which k-mer patterns best represent the observed heterozygosity. We found that the frequency of AAAB k-mers (where only one of the four genomic copies carries a different allele) was greater than or equal to the frequency of AABB k-mers (where two of the genomic copies carry an allele, and the other two copies carry another allele) in nearly all cases (Fig 6).

**Fig 6:**
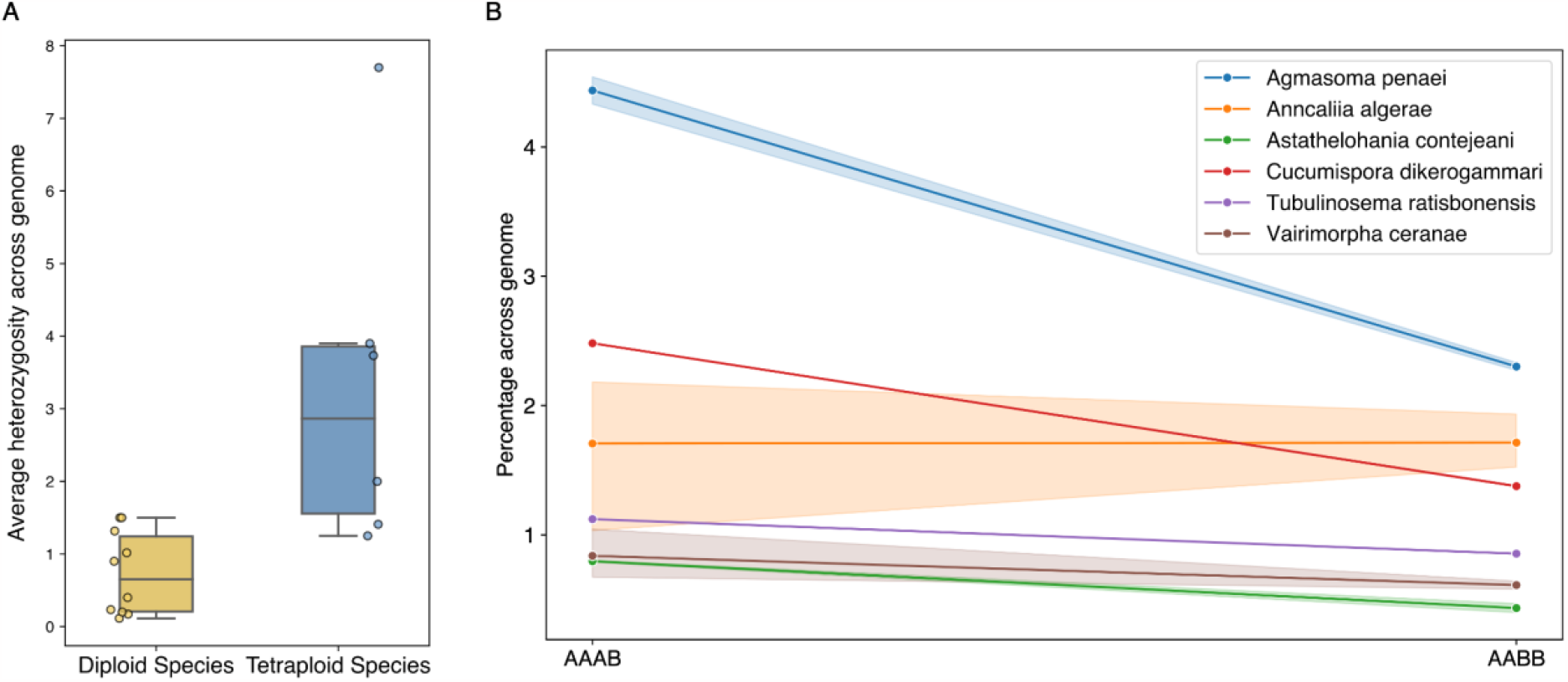
Most diploid and tetraploid genomes are highly homozygous, with the frequency of AAAB k-mers greater than or equal to the frequency of AABB in nearly all cases. (A) Boxplot of mean heterozygosity across the genome per species, calculated using GenomeScope2 heterozygosity estimates (1 - % of AA for diploid species, and 1 - % of AAAA for tetraploid species). (B) Line graph showing mean AAAB and AABB heterozygous k-mer patterns across the genome for each of the six tetraploid species. The shaded region represents the value range observed in the independent samples of each species.

## Discussion

We have used publicly available genomic data to demonstrate that polyploidy is widespread in Microsporidia, with tetraploid species present in three of the six major clades (Nosematida, Glugeida, and Neopereziida [12]), and the orphan lineage represented by *Astahelohania contejeani*.

The presence of polyploidy across the phylum is perplexing, as Microsporidia are obligate intracellular parasites. Such endoparasites have typically been found to have smaller and more compact genomes compared to their free-living counterparts across the tree of life [16,17,21,62]. Indeed, microsporidia have very small genomes, with *Encephalitozoon romaleae* having the smallest eukaryotic genome known, at only 2 Mb [63]. It is thought that the reduction of the genomes of intracellular parasites results from evolutionary pressures to reduce replication time and energy consumption, to maintain a small cell size, and through streamlining due to reliance on host-provided metabolism. Small cell size may be particularly important in parasites which exit their host cells through lysis, as the number of spores they can produce (and thus reproductive potential or R_O_) will be limited by the relative sizes of their cells and the host cell. In line with this, *Nematocida displodere*, which releases its spores through lysis, has a smaller genome than its counterpart *Nematocida parisii*, whose spores are released through exocytosis [20,64]. However our sample size is small, and while spores of the tetraploid species are on average larger than those of the diploids, this difference is not significant (supplementary information Fig S1).

Given the wide range of genome spans in Microsporidia [63,65], it is unclear how strong the selection pressure for small genomes is. If polyploidy arises relatively frequently, and the immediate fitness costs of the larger genome and larger spore size are low, polyploid lineages may arise and persist for short periods of evolutionary time before being driven to extinction. Our limited sampling across Microsporidia precludes examination of the relative persistence of polyploids. Additionally, polyploidy may have a positively selected role in rescuing inbred or asexual microsporidians from accumulated lethal mutations, and in generating allelic diversity in hybrid lineages. There are no clear ecological factors, such as habitats, hosts, and transmission mechanisms that distinguish polyploid species from diploids in our data (supplementary information Figs S2, S3, S4, and S5). While the mechanism of the origin of polyploidy also remains uncertain, it is evident that ploidy is dynamic in Microsporidia.

The majority (>96%) of sites in the polyploid genomes are highly homozygous, with the frequency of AAAB k-mers (where one of the four genomic copies differs from the other three) being greater than or equal to the frequency of AABB k-mers (where the two alternate alleles are equally present) in almost all cases. Auto-polyploidy is *a priori* highly plausible given the intracellular niche of microsporidia and the presumed rarity of partners (for sex as well as for allo-polyploidisation). In an auto-polyploid, we would expect the homeologous chromosomes to show divergence commensurate with the cumulative effect of neutral mutation over time and generations. In the species we assessed as diploid, we identified heterozygosities between 0.5% and 1.5%, whereas in the tetraploids, heterozygosity ranged from 1.2% to 4%. This in itself suggests that the homeologous genome copies in the tetraploids are more different from each other than one would expect from very recent auto-polyploidy. Alternatively, if these tetraploids are recent and derive from allo-polyploidy, the parental genomes are likely to have been as or more divergent than the levels we measured between alleles within the diploids. In both models, residual meiotic recombination between homeologous chromosomes and ongoing inter-homeologue gene conversion would tend to reduce observed divergence between homeologues and thus measured heterozygosity. It is difficult to fully interpret the patterns observed without knowledge of microsporidian mutational and recombination rates, the rate of gene conversion between homeologues, the age of the polyploidisation events, and the frequency of meiosis in tetraploid species’ life cycles. It is of course possible that different tetraploids were created by different routes.

Progress in disentangling the origins and dynamics of polyploidy in Microsporidia is constrained by current short-read whole genome data. Phased, chromosomally assembled genome sequences will be critical in distinguishing between different models, and also in furthering genomic understanding of microsporidian biology. Ongoing reference-quality genome sequencing and genomic surveillance programmes, such as the [Darwin Tree of Life project](https://www.darwintreeoflife.org/) [66] and [BIOSCAN](https://www.sanger.ac.uk/collaboration/bioscan/), often identify microsporidia as cobionts when sequencing host species. These initiatives offer an unrivalled opportunity for the generation of high-quality, phased, long-read microsporidian genomes from a variety of hosts, and will soon allow us to explore questions like the timing of polyploidisation events, their mode, and their genomic impact.

## Data Availability

All the data used in this manuscript is publicly available on NCBI Sequence Read Archive (SRA). Please see supplementary information Tables S1 and S2 for the list of accession numbers.

## Acknowledgments

We would like to thank Dr Jamie Bojko, Dr Claudia Weber, Dr Emmelien Vancaester, and Dr Ellen Cameron for kindly reading the manuscript and commenting on it. We would also like to thank the class of Biodiversity Genomics Academy 2023 workshop “Understanding k-mers and ploidy using Smudgeplot” for recreating our supplementary information table S1 during their practice sessions, demonstrating reproducibility. The authors were kindly supported by Wellcome Trust award 220540/Z/20/A, ‘Wellcome Sanger Institute Quinquennial Review 2021-2026’.

## References

1. Keeling P. Five questions about microsporidia. PLoS Pathog. 2009;5:e1000489.doi:10.1371/journal.ppat.1000489

2. Fokin SI, Di Giuseppe G, Erra F, Dini F. Euplotespora binucleata n. gen., n. sp.(Protozoa: Microsporidia), a parasite infecting the hypotrichous ciliate Euploteswoodruffi, with observations on microsporidian infections in ciliophora. J EukaryotMicrobiol. 2008;55: 214–228. doi:10.1111/j.1550-7408.2008.00322.x

3. Stentiford GD, Feist SW, Stone DM, Bateman KS, Dunn AM. Microsporidia: diverse,dynamic, and emergent pathogens in aquatic systems. Trends Parasitol. 2013;29: 567–578. doi:10.1016/j.pt.2013.08.005

4. Murareanu BM, Sukhdeo R, Qu R, Jiang J, Reinke AW. Generation of a MicrosporidiaSpecies Attribute Database and Analysis of the Extensive Ecological and PhenotypicDiversity of Microsporidia. MBio. 2021;12: e0149021. doi:10.1128/mBio.01490-21

5. Bojko J, Stentiford GD. Microsporidian Pathogens of Aquatic Animals. ExperientiaSuppl. 2022;114: 247–283. doi:10.1007/978-3-030-93306-7_10

6. Nageli C. uber die neue Krankheit der Seidenraupe und verwandte Organismen.[Abstract of report before 33. Versamml. Deutsch. Naturf. u. Aerzte. Bonn, 21 Sept.].Bot Ztg. 1857;15: 760–761. Available: https://cir.nii.ac.jp/crid/1573105974684833920

7. Pasteur L. Etudes sur la maladie des vers à soie: 2.: Notes et documents. Gauthier-Villars; 1870. Available: https://play.google.com/store/books/details?id=y-1rmRQoAa4C

8. Han B, Weiss LM. Microsporidia: Obligate Intracellular Pathogens Within the FungalKingdom. Microbiol Spectr. 2017;5. doi:10.1128/microbiolspec.FUNK-0018-2016

9. Katinka MD, Duprat S, Cornillot E, Méténier G, Thomarat F, Prensier G, et al. Genomesequence and gene compaction of the eukaryote parasite Encephalitozoon cuniculi.Nature. 2001;414: 450–453. doi:10.1038/35106579

10. Heinz E, Williams TA, Nakjang S, Noël CJ, Swan DC, Goldberg AV, et al. The genomeof the obligate intracellular parasite Trachipleistophora hominis: new insights intomicrosporidian genome dynamics and reductive evolution. PLoS Pathog. 2012;8:e1002979. doi:10.1371/journal.ppat.1002979

11. Wadi L, Reinke AW. Evolution of microsporidia: An extremely successful group ofeukaryotic intracellular parasites. PLoS Pathog. 2020;16: e1008276.doi:10.1371/journal.ppat.1008276

12. Bojko J, Reinke AW, Stentiford GD, Williams B, Rogers MSJ, Bass D. Microsporidia: anew taxonomic, evolutionary, and ecological synthesis. Trends Parasitol. 2022;38: 642–659. doi:10.1016/j.pt.2022.05.007

13. Wang L, Li H, Shi W, Qiao Y, Wang P, Yu Z, et al. Whole-genome sequencing andcomparative genomic analysis of a pathogenic Enterocytozoon hepatopenaei strainisolated from Litopenaeus vannamei. Aquac Int. 2023;31: 523–546.doi:10.1007/s10499-022-00990-9

14. Angst P, Pombert J-F, Ebert D, Fields PD. Near chromosome-level genome assemblyof the microsporidium Hamiltosporidium tvaerminnensis. G3. 2023.doi:10.1093/g3journal/jkad185

15. Xiong X, Geden CJ, Bergstralh DT, White RL, Werren JH, Wang X. New insights intothe genome and transmission of the microsporidian pathogen Nosema muscidifuracis.Front Microbiol. 2023;14: 1152586. doi:10.3389/fmicb.2023.1152586

16. Andersson JO, Andersson SG. Insights into the evolutionary process of genomedegradation. Curr Opin Genet Dev. 1999;9: 664–671. doi:10.1016/s0959-437x(99)00024-6

17. Sakharkar KR, Dhar PK, Chow VTK. Genome reduction in prokaryotic obligatoryintracellular parasites of humans: a comparative analysis. Int J Syst Evol Microbiol.2004;54: 1937–1941. doi:10.1099/ijs.0.63090-0

18. Keeling PJ, Corradi N. Shrink it or lose it: balancing loss of function with shrinkinggenomes in the microsporidia. Virulence. 2011;2: 67–70. doi:10.4161/viru.2.1.14606

19. Wiredu Boakye D, Jaroenlak P, Prachumwat A, Williams TA, Bateman KS,Itsathitphaisarn O, et al. Decay of the glycolytic pathway and adaptation to intranuclearparasitism within Enterocytozoonidae microsporidia. Environ Microbiol. 2017;19: 2077–2089. doi:10.1111/1462-2920.13734

20. Jespersen N, Monrroy L, Barandun J. Impact of Genome Reduction in Microsporidia.Experientia Suppl. 2022;114: 1–42. doi:10.1007/978-3-030-93306-7_1

21. Williams BAP, Williams TA, Trew J. Comparative Genomics of Microsporidia.Experientia Suppl. 2022;114: 43–69. doi:10.1007/978-3-030-93306-7_2

22. Burri L, Williams BAP, Bursac D, Lithgow T, Keeling PJ. Microsporidian mitosomesretain elements of the general mitochondrial targeting system. Proceedings of theNational Academy of Sciences. 2006;103: 15916–15920. doi:10.1073/pnas.0604109103

23. Amigó JM, Gracia MP, Salvadó H, Vivarés CP. Pulsed Field Gel Electrophoresis ofThree Microsporidian Parasites of Fish. Acta Protozool. 2002;41: 11–16. Available:https://www.airitilibrary.com/Publication/alDetailedMesh?docid=00651583-200203-201012290004-201012290004-11-16

24. Lee, Heitman, Ironside. Sex and the Microsporidia. Microsporidia: Pathogens of. 2014.Available:https://books.google.com/books?hl=en&lr=&id=k4kZBAAAQBAJ&oi=fnd&pg=PA231&dq=microsporidia+ploidy&ots=oqF0tRPXyc&sig=Ua18f4MX3vgjvbaipK2DYB1v3Kc

25. Lee SC, Corradi N, Byrnes EJ 3rd, Torres-Martinez S, Dietrich FS, Keeling PJ, et al.Microsporidia evolved from ancestral sexual fungi. Curr Biol. 2008;18: 1675–1679.doi:10.1016/j.cub.2008.09.030

26. Banuett F. From dikaryon to diploid. Fungal Biol Rev. 2015;29: 194–208.doi:10.1016/j.fbr.2015.08.001

27. Weidner E. Ultrastructural study of microsporidian development. Zeitschrift fürZellforschung und Mikroskopische Anatomie. 1970;105: 33–54.doi:10.1007/BF00340563

28. Vávra J. Development of the Microsporidia. In: Bulla LA, Cheng TC, editors. Biology ofthe Microsporidia. Boston, MA: Springer US; 1976. pp. 87–109. doi:10.1007/978-1-4684-3114-8_3

29. Maurand J, Vey A. Etudes histopathologique et ultrastructurale de Thelohaniacontejeani (Microsporida, Nosematidae) parasite de l’Ecrevisse Austropotamobiuspallipes Lereboullet. Ann Parasitol Hum Comp. 1973;48: 411–421.doi:10.1051/parasite/1973483411

30. Pretto T, Montesi F, Ghia D, Berton V, Abbadi M, Gastaldelli M, et al. Ultrastructural andmolecular characterization of Vairimorpha austropotamobii sp. nov. (Microsporidia:Burenellidae) and Thelohania contejeani (Microsporidia: Thelohaniidae), two parasitesof the white-clawed crayfish, Austropotamobius pallipes complex (Decapoda:Astacidae). J Invertebr Pathol. 2018;151: 59–75. doi:10.1016/j.jip.2017.11.002

31. Miquel J, Kacem H, Baz-González E, Foronda P, Marchand B. Ultrastructural andmolecular study of the microsporidian Toguebayea baccigeri n. gen., n. sp., ahyperparasite of the digenean trematode Bacciger israelensis (Faustulidae), a parasiteof Boops boops (Teleostei, Sparidae). Parasite. 2022;29: 2.doi:10.1051/parasite/2022007

32. Sprague V, Vernick SH. The ultrastructure of Encephalitozoon cuniculi (Microsporida,Nosematidae) and its taxonomic significance. J Protozool. 1971;18: 560–569.doi:10.1111/j.1550-7408.1971.tb03376.x

33. Desportes I. ULTRASTRUCTURE DE STEMPELLIA MUTABILIS LEGER ET HESSE,MICROSPORIDIE PARASITE DE L’EPHEMERE EPHEMERA VULGATA L. 1976 [cited29 Dec 2022]. Available: https://pascal-francis.inist.fr/vibad/index.php?action=getRecordDetail&idt=PASCAL7750051096

34. Loubès C, Maurand J, Rousset-Galangau V. Presence of synaptonematic complexes inthe biological cycle of Gurleya chironomi Loubes and Maurand, 1975: an argument infavor of sexuality in microsporidia. CR Hebd Seances Acad Sci Ser D Sci Nat.1976;282: 1025–1027. Available: https://europepmc.org/article/med/821628

35. Loubès C. [Meiosis in Microsporidia: effects on biological cycles]. J Protozool. 1979;26:200–208. doi:10.1111/j.1550-7408.1979.tb02761.x

36. Vivares CP, Sprague V. The fine structure of Ameson pulvis (Microspora, Microsporida)and its implications regarding classification and chromosome cycle. J Invertebr Pathol.1979;33: 40–52. doi:10.1016/0022-2011(79)90128-9

37. Hazard EI, Andreadis TG, Joslyn DJ, Ellis EA. Meiosis and Its Implications in the LifeCycles of Amblyospora and Parathelohania (Microspora). J Parasitol. 1979;65: 117–122. doi:10.2307/3280215

38. Hazard EI, Brookbank JW. Karyogamy and meiosis in an Amblyospora sp. (Microspora)in the mosquito Culex salinarius. J Invertebr Pathol. 1984;44: 3–11. doi:10.1016/0022-2011(84)90039-9

39. Hazard EI, Fukuda T, Becnel JJ. Life cycle ofCulicosporella lunata(hazard & savage,1970) Weiser, 1977 (Microspora) as revealed in the light microscope with aredescription of the genus and Species1. J Protozool. 1984;31: 385–391.doi:10.1111/j.1550-7408.1984.tb02984.x

40. Becnel JJ, Hazard EI, Fukuda T, Sprague V. Life cycle ofCulicospora magna(Kudo,1920) (microsporida: Culicosporidae) inCulex restuansTheobald with special referenceto Sexuality1. J Protozool. 1987;34: 313–322. doi:10.1111/j.1550-7408.1987.tb03182.x

41. Canning EU. Nuclear division and chromosome cycle in microsporidia. Biosystems.1988;21: 333–340. doi:10.1016/0303-2647(88)90030-5

42. Becnel JJ, Sprague V, Fukuda T, Hazard EI. Development of Edhazardia aedis (Kudo,1930) n. g., n. comb. (Microsporida: Amblyosporidae) in the mosquito Aedes aegypti (L.)(Diptera: Culicidae). J Protozool. 1989;36: 119–130. doi:10.1111/j.1550-7408.1989.tb01057.x

43. Becnel JJ. Horizontal transmission and subsequent development of Amblyosporacalifornica (Microsporida: Amblyosporidae) in the intermediate and definitive hosts. DisAquat Organ. 1992;13: 17–28. Available: https://www.int-res.com/articles/dao/13/d013p017.pdf

44. Sokolova YY, Fuxa JR. Biology and life-cycle of the microsporidium Kneallhaziasolenopsae Knell Allan Hazard 1977 gen. n., comb. n., from the fire ant Solenopsisinvicta. Parasitology. 2008;135: 903–929. doi:10.1017/S003118200800440X

45. Cali A. Morphogenesis in the genus Nosema. ProcIVth IntColloqInsect Pathol. 1971;4:104–112. Available: https://ci.nii.ac.jp/naid/10004755181/

46. Youssef NN, Hammond DM. The fine structure of the developmental stages of themicrosporidian Nosema apis Zander. Tissue Cell. 1971;3: 283–294. doi:10.1016/s0040-8166(71)80023-x

47. Cuomo CA, Desjardins CA, Bakowski MA, Goldberg J, Ma AT, Becnel JJ, et al.Microsporidian genome analysis reveals evolutionary strategies for obligate intracellulargrowth. Genome Res. 2012;22: 2478–2488. doi:10.1101/gr.142802.112

48. Pombert J-F, Xu J, Smith DR, Heiman D, Young S, Cuomo CA, et al. Complete genomesequences from three genetically distinct strains reveal high intraspecies geneticdiversity in the microsporidian Encephalitozoon cuniculi. Eukaryot Cell. 2013;12: 503–511. doi:10.1128/EC.00312-12

49. Selman M, Sak B, Kváč M, Farinelli L, Weiss LM, Corradi N. Extremely reduced levelsof heterozygosity in the vertebrate pathogen Encephalitozoon cuniculi. Eukaryot Cell.2013;12: 496–502. doi:10.1128/EC.00307-12

50. Pelin A, Moteshareie H, Sak B, Selman M, Naor A, Eyahpaise M-È, et al. The genomeof an Encephalitozoon cuniculi type III strain reveals insights into the genetic diversityand mode of reproduction of a ubiquitous vertebrate pathogen. Heredity. 2016;116:458–465. doi:10.1038/hdy.2016.4

51. Pelin A, Selman M, Aris-Brosou S, Farinelli L, Corradi N. Genome analyses suggest thepresence of polyploidy and recent human-driven expansions in eight global populationsof the honeybee pathogen Nosema ceranae. Environ Microbiol. 2015;17: 4443–4458.doi:10.1111/1462-2920.12883

52. Bernander R, Palm JE, Svärd SG. Genome ploidy in different stages of the Giardialamblia life cycle. Cell Microbiol. 2001;3: 55–62. doi:10.1046/j.1462-5822.2001.00094.x

53. Leinonen R, Sugawara H, Shumway M, International Nucleotide Sequence DatabaseCollaboration. The sequence read archive. Nucleic Acids Res. 2011;39: D19–21.doi:10.1093/nar/gkq1019

54. Marçais G, Kingsford C. A fast, lock-free approach for efficient parallel counting ofoccurrences of k-mers. Bioinformatics. 2011;27: 764–770.doi:10.1093/bioinformatics/btr011

55. Ranallo-Benavidez TR, Jaron KS, Schatz MC. GenomeScope 2.0 and Smudgeplot for reference-free profiling of polyploid genomes. Nat Commun. 2020;11: 1–10.doi:10.1038/s41467-020-14998-3

56. Miller CS, Baker BJ, Thomas BC, Singer SW, Banfield JF. EMIRGE: reconstruction of full-length ribosomal genes from microbial community short read sequencing data.Genome Biol. 2011;12: R44. doi:10.1186/gb-2011-12-5-r44

57. Gruber-Vodicka HR, Seah BKB, Pruesse E. phyloFlash: Rapid Small-Subunit rRNA Profiling and Targeted Assembly from Metagenomes. mSystems. 2020;5.doi:10.1128/mSystems.00920-20

58. NCBI Resource Coordinators. Database resources of the National Center for Biotechnology Information. Nucleic Acids Res. 2016;44: D7–19.doi:10.1093/nar/gkv1290

59. Katoh K, Misawa K, Kuma K, Miyata T. MAFFT: a novel method for rapid multiple sequence alignment based on fast Fourier transform. Nucleic Acids Res. 2002;30:3059–3066. doi:10.1093/nar/gkf436

60. Minh BQ, Schmidt HA, Chernomor O, Schrempf D, Woodhams MD, von Haeseler A, et al. IQ-TREE 2: New Models and Efficient Methods for Phylogenetic Inference in the Genomic Era. Mol Biol Evol. 2020;37: 1530–1534. doi:10.1093/molbev/msaa015

61. James TY, Pelin A, Bonen L, Ahrendt S, Sain D, Corradi N, et al. Shared Signatures of Parasitism and Phylogenomics Unite Cryptomycota and Microsporidia. Curr Biol.2013;23: 1548–1553. doi:10.1016/j.cub.2013.06.057

62. Žárský V, Karnkowska A, Boscaro V, Trznadel M, Whelan TA, Hiltunen-Thorén M, et al. Contrasting outcomes of genome reduction in mikrocytids and microsporidians. BMCBiol. 2023;21: 137. doi:10.1186/s12915-023-01635-w

63. Pombert J-F, Selman M, Burki F, Bardell FT, Farinelli L, Solter LF, et al. Gain and loss of multiple functionally related, horizontally transferred genes in the reduced genomes of two microsporidian parasites. Proc Natl Acad Sci U S A. 2012;109: 12638–12643.doi:10.1073/pnas.1205020109

64. Zhang G, Sachse M, Prevost M-C, Luallen RJ, Troemel ER, Félix M-A. A Large Collection of Novel Nematode-Infecting Microsporidia and Their Diverse Interactions with Caenorhabditis elegans and Other Related Nematodes. PLoS Pathog. 2016;12:e1006093. doi:10.1371/journal.ppat.1006093

65. Desjardins CA, Sanscrainte ND, Goldberg JM, Heiman D, Young S, Zeng Q, et al.Contrasting host–pathogen interactions and genome evolution in two generalist and specialist microsporidian pathogens of mosquitoes. Nat Commun. 2015;6: 1–12.doi:10.1038/ncomms8121

66. The Darwin Tree of Life Project Consortium, Blaxter M, Mieszkowska N, Palma FD,Holland P, Durbin R, et al. Sequence locally, think globally: The Darwin Tree of Life Project. Proceedings of the National Academy of Sciences. 2022;119: e2115642118.doi:10.1073/pnas.2115642118

