## Supplementary Table 1 for "Polyploidy is widespread in Microsporidia"

Table S1: SRA accession numbers, species name, linear GenomeScope2 plots, transformed GenomeScope2 plots, and Smudgeplot plots of the 66 samples for which we were able to estimate ploidy in this study.

| Index | SRA Accession | Species | Estimated Ploidy | Genomscope2 Linear Plot | Genomscope2 Transformed Linear Plot | Smudgeplot | Comments |
|---|---|---|---|---|---|---|---|
| 1 | SRR17317295 | Vairimorpha ceranae | 4 | 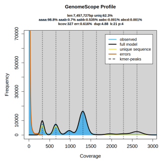 | 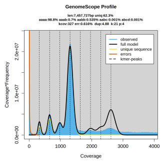 | 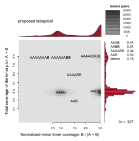 | |
| 2 | SRR17317294 | Vairimorpha ceranae | 4 | 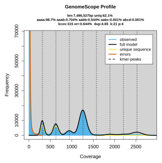 | 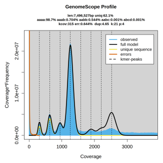 | 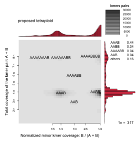 | |
| 3 | SRR17317293 | Vairimorpha ceranae | 4 | 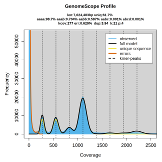 | 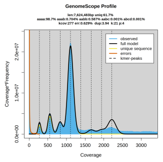 | 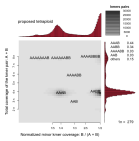 | |
| 4 | SRR17317292 | Vairimorpha ceranae | 4 | 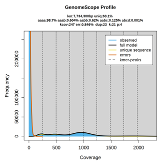 | 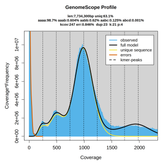 | 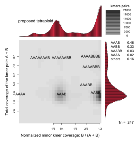 | |
| 5 | SRR17317291 | Vairimorpha ceranae | 4 | 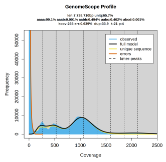 | 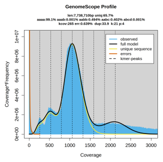 | 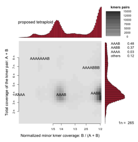 | |
| 6 | SRR17317296 | Vairimorpha ceranae | 4 | 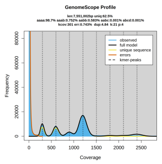 | 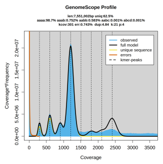 | 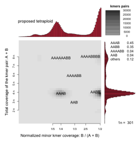 | |
| 7 | SRR18590839 | Vairimorpha ceranae | 4 | 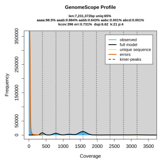 | 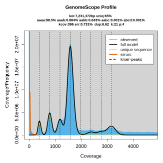 | 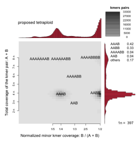 | |
| 8 | SRR18590838 | Vairimorpha ceranae | 4 | 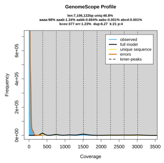 | 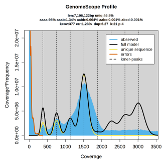 | 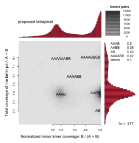 | |
| 9 | SRR18590835 | Vairimorpha ceranae | 4 | 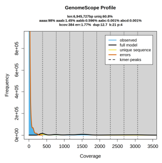 | 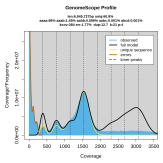 | 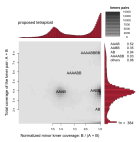 | Low coverage contaminant |

| Index | SRA Accession | Species | Estimated Ploidy | Genomscope2 Linear Plot | Genomscope2 Transformed Linear Plot | Smudgeplot | Comments |
|---|---|---|---|---|---|---|---|
| 10 | SRR23560257 | Encephalitozoon hellem | 2 | 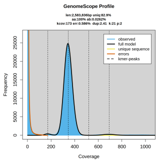 | 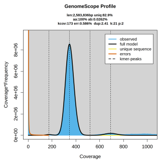 | 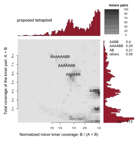 | Genome is highly homozygous. The 4n smudge appears because imperfect duplications dominate over nearly negligible heterozygosity |
| 11 | SRR23560258 | Encephalitozoon hellem | 2 | 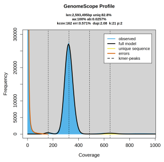 | 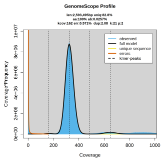 | 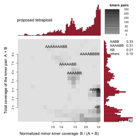 | Genome is highly homozygous. The 4n smudge appears because imperfect duplications dominate over nearly negligible heterozygosity |
| 12 | SRR23560420 | Encephalitozoon hellem | 2 | 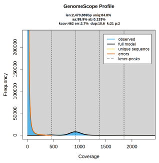 | 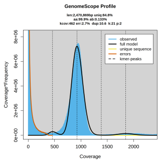 | 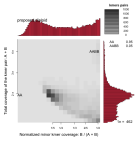 | Genome is highly homozygous. Signal at the edges of the smudgeplot is noise |
| 13 | SRR24007516 | Encephalitozoon intestinalis | 2 | 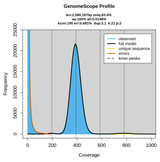 | 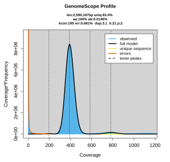 | 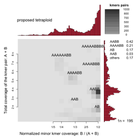 | Genome is highly homozygous. The 4n smudge appears because imperfect duplications dominate over nearly negligible heterozygosity |
| 14 | SRR24007515 | Encephalitozoon intestinalis | 2 | 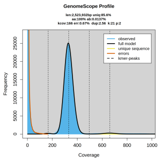 | 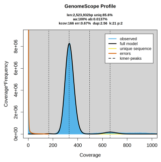 | 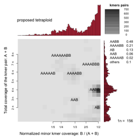 | Genome is highly homozygous. The 4n smudge appears because |

| Index | SRA Accession | Species | Estimated Ploidy | Genomscope2 Linear Plot | Genomscope2 Transformed Linear Plot | Smudgeplot | Comments |
|---|---|---|---|---|---|---|---|
| | | | | | | | imperfect duplications dominate over nearly negligible heterozygosity |
| 15 | SRR24007514 | Encephalitozoon intestinalis | 2 | 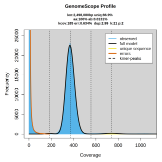 | 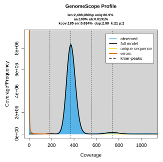 | 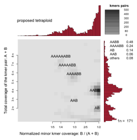 | Genome is highly homozygous. The 4n smudge appears because imperfect duplications dominate over nearly negligible heterozygosity |
| 16 | SRR16954902 | Hamiltosporidium magnivora | 2 | 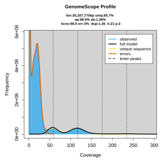 | 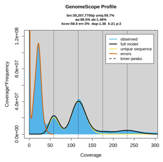 | 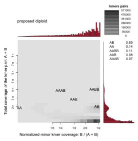 | Low coverage contaminant |
| 17 | SRR16954901 | Hamiltosporidium tvaerminnensis | 2 | 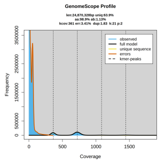 | 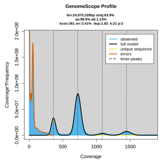 | 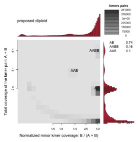 | Low coverage contaminant |
| 18 | SRR16954899 | Hamiltosporidium tvaerminnensis | 2 | 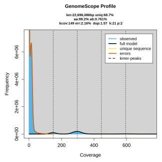 | 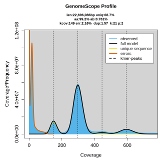 | 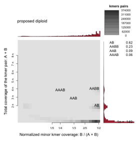 | Low coverage contaminant |
| 19 | SRR16954898 | Hamiltosporidium tvaerminnensis | 2 | 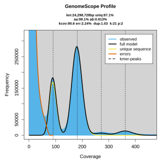 | 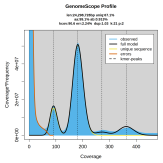 | 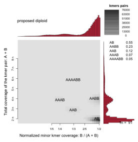 | |
| 20 | SRR16954897 | Hamiltosporidium tvaerminnensis | 2 | 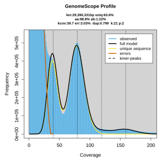 | 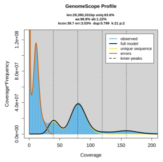 | 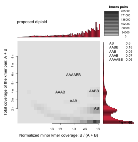 | Low coverage contaminant |
| 21 | SRR16954895 | Hamiltosporidium tvaerminnensis | 2 | 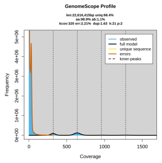 | 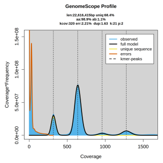 | 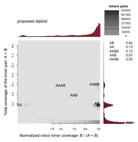 | Low coverage contaminant |

| Index | SRA Accession | Species | Estimated Ploidy | Genomscope2 Linear Plot | Genomscope2 Transformed Linear Plot | Smudgeplot | Comments |
|---|---|---|---|---|---|---|---|
| 22 | SRR16954906 | Hamiltosporidium tvaerminnensis | 2 | 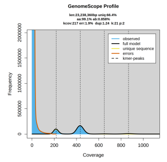 | 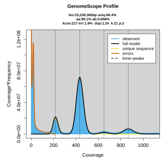 | 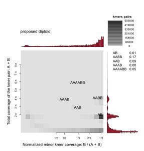 | Low coverage contaminant |
| 23 | SRR16954905 | Hamiltosporidium tvaerminnensis | 2 | 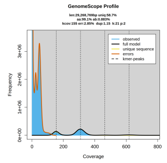 | 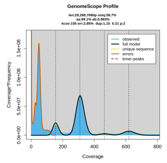 | 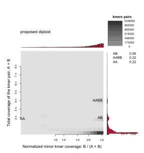 | Low coverage contaminant |
| 24 | SRR16954904 | Hamiltosporidium tvaerminnensis | 2 | 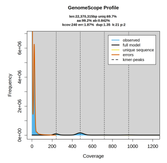 | 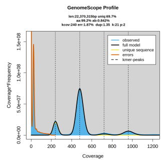 | 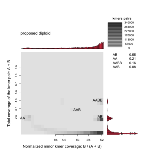 | Low coverage contaminant |
| 25 | SRR23214363 | Nematocida ausubeli | 2 | 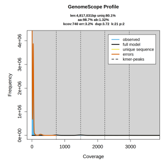 | 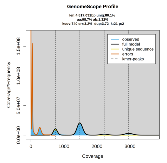 | 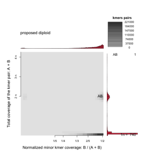 | Low coverage contaminant |
| 26 | SRR23214355 | Nematocida ausubeli | 2 | 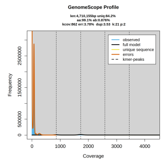 | 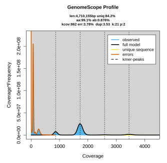 | 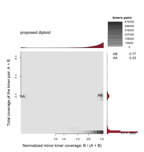 | Sample has a low coverage contaminant |
| 27 | SRR23214353 | Nematocida ausubeli | 2 | 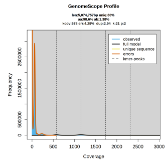 | 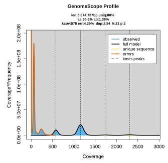 | 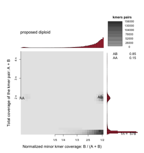 | Low coverage contaminant |
| 28 | SRR23214358 | Nematocida ausubeli | 2 | 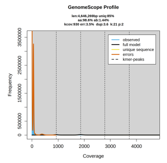 | 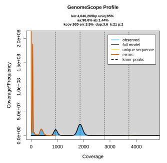 | 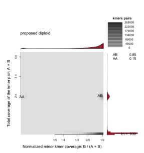 | Low coverage contaminant |
| 29 | SRR17622377 | Nematocida major | 2 | 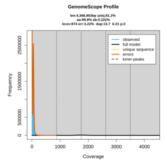 | 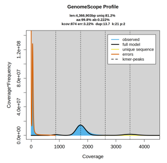 | 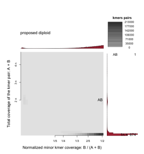 | Low coverage contaminant |
| 30 | SRR23214350 | Pancytospora philotis | 2 | 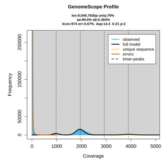 | 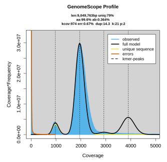 | 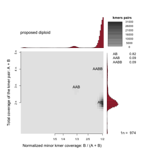 | |
| 31 | ERR3154977 | Tubulinosema ratisbonensis | 4 | 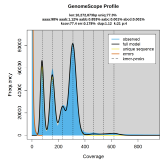 | 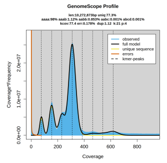 | 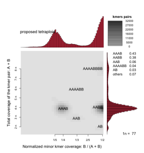 | |

| Index | SRA Accession | Species | Estimated Ploidy | Genomscope2 Linear Plot | Genomscope2 Transformed Linear Plot | Smudgeplot | Comments |
|---|---|---|---|---|---|---|---|
| 32 | SRR2105612 | Encephalitozoon cuniculi EcunIII-L | 2 | 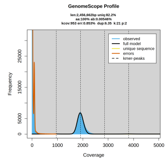 | 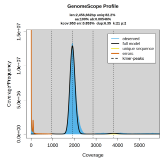 | 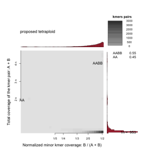 | |
| 33 | SRR8476225 | Astathelohania contejeani | 4 | 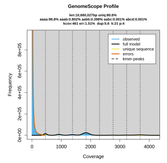 | 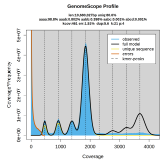 | 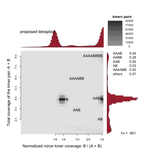 | |
| 34 | SRR8476226 | Astathelohania contejeani | 4 | 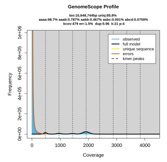 | 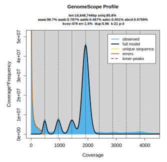 | 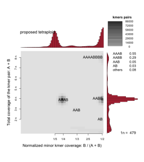 | |
| 35 | SRR8495097 | Cucumispora dikerogammari | 4 | 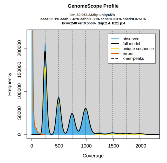 | 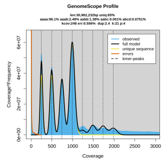 | 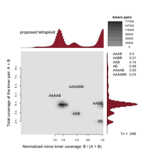 | |
| 36 | SRR8495098 | Cucumispora dikerogammari | 4 | 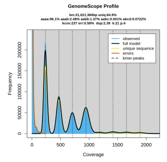 | 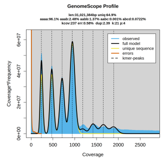 | 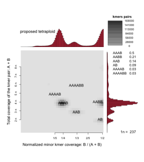 | |
| 37 | SRR9597065 | Hamiltosporidium tvaerminnensis | 2 | 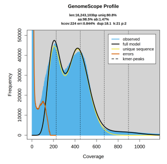 | 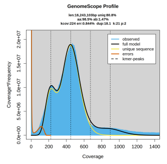 | 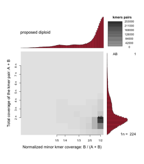 | |
| 38 | SRR14017862 | Encephalitozoon hellem | 2 | 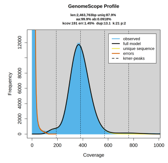 | 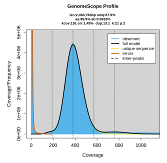 | 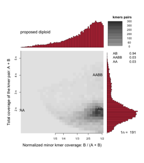 | |
| 39 | SRR14062087 | Encephalitozoon hellem | 2 | 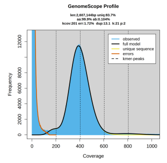 | 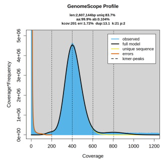 | 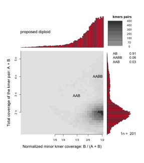 | |
| 40 | SRR16954910 | Hamiltosporidium tvaerminnensis | 2 | 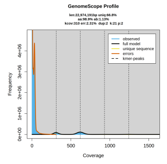 | 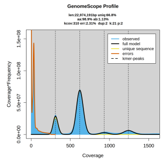 | 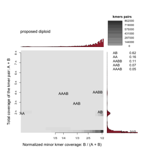 | Low coverage contaminant |
| 41 | SRR16954909 | Hamiltosporidium tvaerminnensis | 2 | 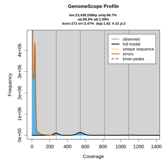 | 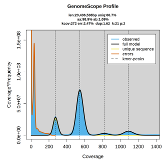 | 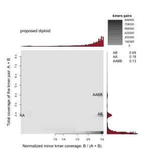 | Low coverage contaminant |

| Index | SRA Accession | Species | Estimated Ploidy | Genomscope2 Linear Plot | Genomscope2 Transformed Linear Plot | Smudgeplot | Comments |
|---|---|---|---|---|---|---|---|
| 42 | SRR16954908 | Hamiltosporidium tvaerminnensis | 2 | 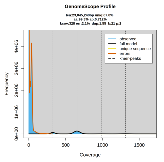 | 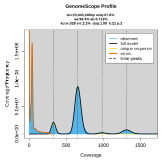 | 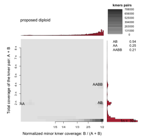 | Low coverage contaminant |
| 43 | SRR543737 | Anncaliia algerae | 4 | 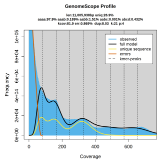 | 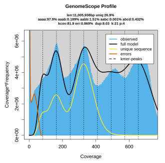 | 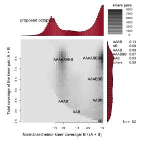 | |
| 44 | SRR630040 | Anncaliia algerae | 4 | 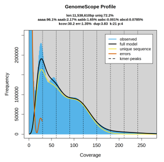 | 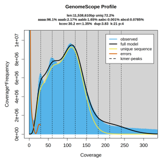 | 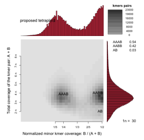 | |
| 45 | SRR489790 | Anncaliia algerae | 4 | 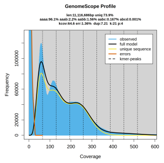 | 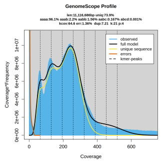 | 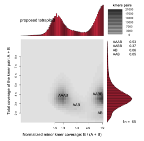 | |
| 46 | SRR489792 | Anncaliia algerae | 4 | 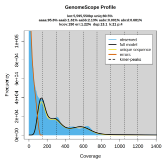 | 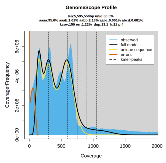 | 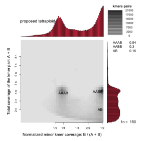 | |
| 47 | SRR653671 | Anncaliia algerae PRA109 | 4 | 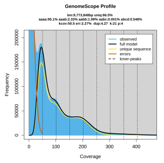 | 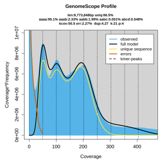 | 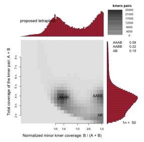 | |
| 48 | SRR122310 | Encephalitozoon cuniculi | 2 | 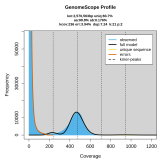 | 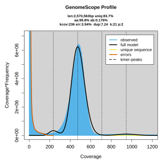 | 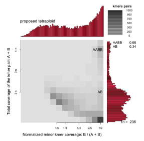 | |
| 49 | SRR122316 | Encephalitozoon cuniculi | 2 | 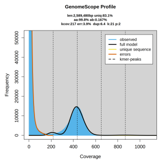 | 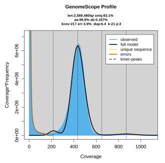 | 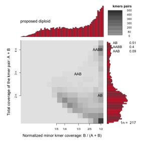 | |
| 50 | SRR122317 | Encephalitozoon cuniculi | 2 | 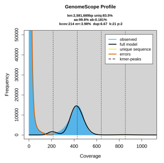 | 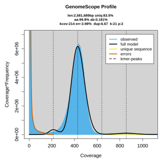 | 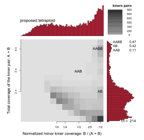 | |
| 51 | SRR065293 | Vittaforma corneae | 2 | 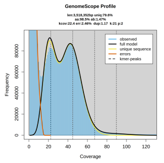 | 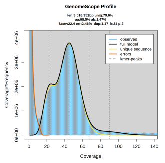 | 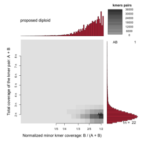 | |

| Index | SRA Accession | Species | Estimated Ploidy | Genomscope2 Linear Plot | Genomscope2 Transformed Linear Plot | Smudgeplot | Comments |
|---|---|---|---|---|---|---|---|
| 52 | SRR926312 | Pseudoloma neurophilia | 2 | 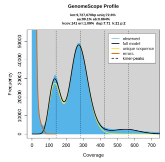 | 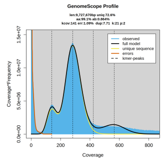 | 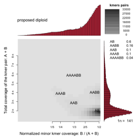 | |
| 53 | SRR926320 | Vairimorpha ceranae | 4 | 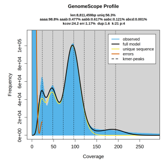 | 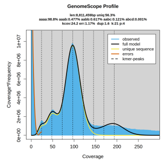 | 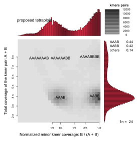 | |
| 54 | SRR1596197 | Vairimorpha ceranae | 4 | 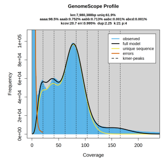 | 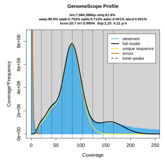 | 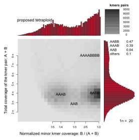 | |
| 55 | SRR926341 | Agmasoma penaei 2012 - Plaquemines Parish LA | 4 | 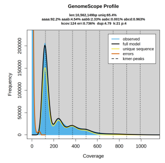 | 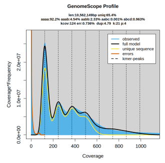 | 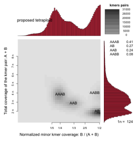 | |
| 56 | SRR926400 | Agmasoma penaei 2012 - Plaquemines Parish LA | 4 | 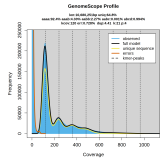 | 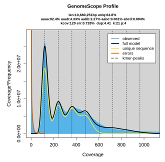 | 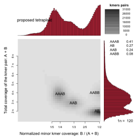 | |
| 57 | SRR058692 | Nematocida ausubeli | 2 | 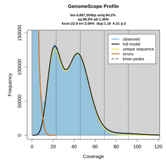 | 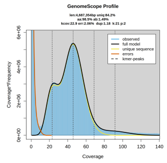 | 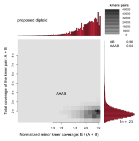 | |
| 58 | SRR350188 | Nematocida ausubeli | 2 | 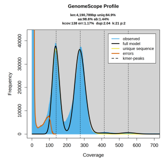 | 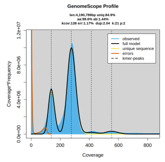 | 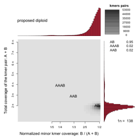 | |
| 59 | SRR17853474 | Encephalitozoon hellem | 2 | 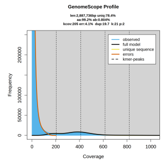 | 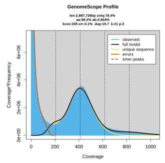 | 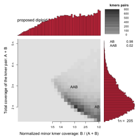 | Smudgeplot is noisy, but genome appears to be highly homozygous |
| 60 | SRR17853475 | Encephalitozoon hellem | 2 | 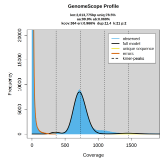 | 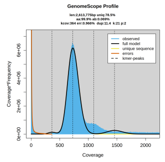 | 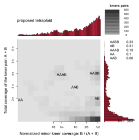 | Smudgeplot is noisy, but genome appears to be highly homozygous |

| Index | SRA Accession | Species | Estimated Ploidy | Genomscope2 Linear Plot | Genomscope2 Transformed Linear Plot | Smudgeplot | Comments |
|---|---|---|---|---|---|---|---|
| 61 | SRR17865589 | Encephalitozoon intestinalis | 2 | 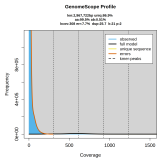 | 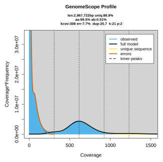 | 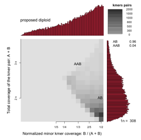 | Smudgeplot is noisy, but genome appears to be highly homozygous |
| 62 | SRR17865590 | Encephalitozoon intestinalis | 2 | 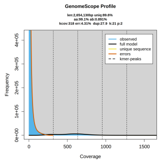 | 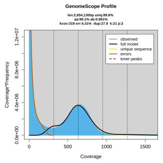 | 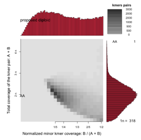 | Smudgeplot is noisy, but genome appears to be highly homozygous |
| 63 | SRR17865591 | Encephalitozoon intestinalis | 2 | 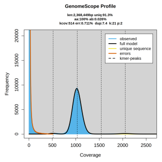 | 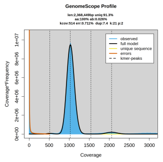 | 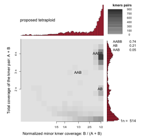 | Smudgeplot is noisy, but genome appears to be highly homozygous. The reason for a smudge at 4n is that due to the high homozygosity, imperfect duplications appear overrepresented. |
| 64 | SRR17858634 | Encephalitozoon cuniculi | 2 | 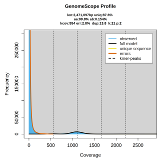 | 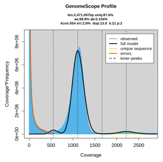 | 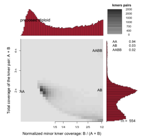 | Smudgeplot is really noisy, but genome appears to be highly homozygous |
| 65 | SRR17858635 | Encephalitozoon cuniculi | 2 | 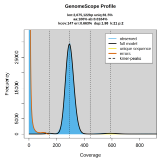 | 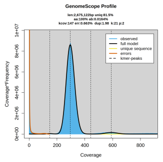 | 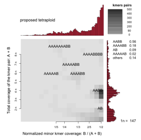 | Smudgeplot is noisy, but genome appears to be highly homozygous. The reason for a smudge at 4n is that due to the high homozygosity, imperfect duplications appear overrepresented. |

| Index | SRA Accession | Species | Estimated Ploidy | Genomscope2 Linear Plot | Genomscope2 Transformed Linear Plot | Smudgeplot | Comments |
|---|---|---|---|---|---|---|---|
| 66 | SRR17858636 | Encephalitozoon cuniculi | 2 | 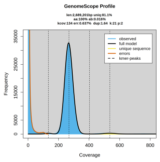 | 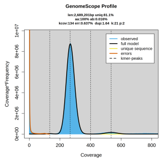 | 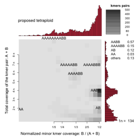 | Smudgeplot is noisy, but genome appears to be highly homozygous. The reason for a smudge at 4n is that due to the high homozygosity, imperfect duplications appear overrepresented. |
