## Supplementary Information for "Polyploidy is widespread in Microsporidia"

**ORCID**

| | |
|---|---|
| Amjad Khalaf | 0000-0003-1297-1181 |
| Kamil S. Jaron | 0000-0003-1470-5450 |
| Mark L. Blaxter | 0000-0003-2861-949X |
| Mara K. N. Lawniczak | 0000-0002-3006-2080 |

This PDF file includes:

- Supplementary Table 2 to Supplementary Table 4
- Supplementary Figure 1 to Supplementary Figure 5
- Supplementary Text 1

Supplied as separate files:

- Supplementary Table 1

**Table S2: SRA accession numbers, species name, and reason for exclusion of the SRA samples for which we were not able to reliably estimate ploidy in this study.**

| Accession | Species | Reason for Filtering Out |
| --- | --- | --- |
| SRR17317290 | Vairimorpha ceranae | Contaminated sample |
| SRR18590837 | Vairimorpha ceranae | Smudgeplot pattern unclear, cannot be interpreted with current data. |
| SRR24032065 | Alternosema sp. JBojko-2023a | No evident peaks |
| SRR18339014 | Anncaliia algerae | Low coverage |
| SRR18590836 | Vairimorpha ceranae | Smudgeplot pattern unclear, cannot be interpreted with current data. |
| SRR23214359 | Enteropsectra breve | Smudgeplot pattern unclear, cannot be interpreted with current data. |
| SRR16954900 | Hamiltosporidium tvaerminnensis | Sample dominated by host |
| SRR16954907 | Hamiltosporidium tvaerminnensis | Contaminated sample |
| SRR18339013 | Anncaliia algerae | No evident peaks |
| SRR23214362 | Nematocida ausubeli | Low coverage |
| SRR23214354 | Nematocida homosporus | Contaminated sample |
| SRR23214366 | Nematocida minor | Contaminated sample |
| SRR23214357 | Nematocida parisii | Contaminated sample |
| SRR23214356 | Nematocida parisii | Contaminated sample |
| SRR23214352 | Nematocida parisii | Contaminated sample |
| SRR23214351 | Nematocida parisii | Contaminated sample |
| SRR23214361 | Nematocida parisii | Contaminated sample |
| SRR23214360 | Nematocida parisii | Contaminated sample |
| SRR23214365 | Nematocida sp. AWRm77 | Contaminated sample |
| SRR23214349 | Nematocida sp. AWRm78 | Contaminated sample |
| SRR23214364 | Nematocida sp. AWRm79 | Contaminated sample |
| SRR23214348 | Nematocida sp. AWRm80 | Contaminated sample |
| SRR23214347 | Nematocida sp. LUAm1 | Contaminated sample |
| SRR23214346 | Nematocida sp. LUAm2 | Contaminated sample |
| SRR23214345 | Nematocida sp. LUAm3 | Contaminated sample |
| SRR23214344 | Pancytospora epiphaga | Contaminated sample |

| | | |
|---|---|---|
| ERR3155584 | Tubulinosema ratisbonensis | No evident peaks |
| ERR3155585 | Tubulinosema ratisbonensis | No evident peaks |
| SRR926331 | Vairimorpha ceranae | Low coverage |
| SRR926332 | Vairimorpha ceranae | Low coverage |
| SRR926321 | Vairimorpha ceranae | Contaminated sample |
| SRR3673305 | Nosema pieriae | No evident peaks |
| SRR7178080 | Vairimorpha ceranae | Contaminated sample |
| SRR7178077 | Vairimorpha ceranae | Contaminated sample |
| SRR7178078 | Vairimorpha ceranae | Contaminated sample |
| SRR7178079 | Vairimorpha ceranae | Contaminated sample |
| SRR8481863 | Nosema granulosis | No evident peaks |
| SRR8481864 | Nosema granulosis | No evident peaks |
| SRR8481865 | Nosema granulosis | No evident peaks |
| SRR8481866 | Nosema granulosis | No evident peaks |
| SRR8481867 | Nosema granulosis | No evident peaks |
| SRR8481868 | Nosema granulosis | No evident peaks |
| SRR8481869 | Nosema granulosis | No evident peaks |
| SRR8481870 | Nosema granulosis | No evident peaks |
| SRR8481871 | Nosema granulosis | No evident peaks |
| SRR8481872 | Nosema granulosis | No evident peaks |
| SRR8481873 | Nosema granulosis | No evident peaks |
| SRR8481874 | Nosema granulosis | No evident peaks |
| SRR8481875 | Nosema granulosis | No evident peaks |
| SRR8481876 | Nosema granulosis | No evident peaks |
| SRR8481877 | Nosema granulosis | No evident peaks |
| SRR8481878 | Nosema granulosis | No evident peaks |
| SRR8494486 | Dictyocoela roeselum | No evident peaks |
| SRR8494487 | Dictyocoela roeselum | No evident peaks |
| SRR8494488 | Dictyocoela roeselum | No evident peaks |
| SRR8494489 | Dictyocoela roeselum | No evident peaks |
| SRR8494490 | Dictyocoela muelleri | No evident peaks |
| SRR8494491 | Dictyocoela muelleri | No evident peaks |
| SRR8494492 | Dictyocoela muelleri | No evident peaks |
| SRR8494493 | Dictyocoela muelleri | No evident peaks |
| SRR8536193 | Vairimorpha ceranae | No evident peaks |

| | | |
|---|---|---|
| SRR9597067 | Hamiltosporidium tvaerminnensis | Low coverage |
| SRR9597064 | Hamiltosporidium magnivora | Low coverage |
| SRR9597066 | Hamiltosporidium magnivora | Low coverage |
| SRR9597070 | Ordospora colligata | Sample dominated by host |
| SRR9597069 | Ordospora colligata | Sample dominated by host |
| SRR9597068 | Ordospora colligata | Sample dominated by host |
| SRR10339155 | Enterospora | No evident peaks |
| SRR10948363 | Vairimorpha ceranae | Contaminated sample |
| SRR10948362 | Vairimorpha ceranae | Contaminated sample |
| SRR12486971 | Anncaliia algerae | Low coverage |
| SRR530631 | Anncaliia algerae PRA109 | Low coverage |
| SRR122343 | Anncaliia algerae PRA109 | No evident peaks |
| SRR202953 | Anncaliia algerae PRA109 | No evident peaks |
| SRR202954 | Anncaliia algerae PRA109 | No evident peaks |
| SRR653432 | Anncaliia algerae PRA109 | Low coverage |
| SRR653669 | Anncaliia algerae PRA109 | Contaminated sample |
| SRR653670 | Anncaliia algerae PRA109 | Contaminated sample |
| SRR653429 | Anncaliia algerae PRA109 | Low coverage |
| SRR052647 | Anncaliia algerae PRA109 | No evident peaks |
| SRR057726 | Anncaliia algerae PRA109 | No evident peaks |
| SRR057727 | Anncaliia algerae PRA109 | No evident peaks |
| SRR122312 | Encephalitozoon cuniculi | Contaminated sample |
| SRR122313 | Encephalitozoon cuniculi | Contaminated sample |
| SRR122314 | Encephalitozoon cuniculi | Contaminated sample |
| SRR122315 | Encephalitozoon cuniculi | Low coverage |
| SRR122309 | Encephalitozoon cuniculi | Contaminated sample |
| SRR122311 | Encephalitozoon cuniculi | Contaminated sample |
| SRR064420 | Edhazardia aedis | No evident peaks |
| SRR064202 | Edhazardia aedis | No evident peaks |
| SRR064203 | Edhazardia aedis | No evident peaks |
| SRR064416 | Edhazardia aedis | No evident peaks |
| SRR070507 | Edhazardia aedis | No evident peaks |
| SRR070508 | Edhazardia aedis | No evident peaks |
| SRR070509 | Edhazardia aedis | No evident peaks |

| | | |
|---|---|---|
| SRR070510 | Edhazardia aedis | No evident peaks |
| SRR070511 | Edhazardia aedis | No evident peaks |
| SRR070512 | Edhazardia aedis | No evident peaks |
| SRR070514 | Edhazardia aedis | No evident peaks |
| SRR071362 | Edhazardia aedis | No evident peaks |
| SRR071363 | Edhazardia aedis | No evident peaks |
| SRR071364 | Edhazardia aedis | No evident peaks |
| SRR071365 | Edhazardia aedis | No evident peaks |
| SRR052646 | Vavraia culicis | No evident peaks |
| SRR062126 | Vavraia culicis | No evident peaks |
| SRR064422 | Vavraia culicis | No evident peaks |
| SRR122342 | Vavraia culicis | No evident peaks |
| SRR064421 | Nematocida ausubeli | No evident peaks |
| SRR068584 | Anncaliia algerae | No evident peaks |
| SRR068585 | Anncaliia algerae | No evident peaks |
| SRR068586 | Anncaliia algerae | No evident peaks |
| SRR068587 | Anncaliia algerae | No evident peaks |
| SRR058694 | Vittaforma corneae | Low coverage |
| SRR058934 | Vittaforma corneae | Low coverage |
| SRR057728 | Vittaforma corneae | Low coverage |
| SRR065289 | Vittaforma corneae | No evident peaks |
| SRR064417 | Vittaforma corneae | Low coverage |
| SRR064418 | Vittaforma corneae | No evident peaks |
| SRR064419 | Vittaforma corneae | No evident peaks |
| SRR070313 | Vittaforma corneae | Low coverage |
| SRR070314 | Vittaforma corneae | No evident peaks |
| SRR062125 | Nematocida ausubeli | Low coverage |
| SRR636827 | Encephalitozoon cuniculi | Low coverage |
| SRR765863 | Nosema apis BRL 01 | No evident peaks |
| SRR765864 | Nosema apis BRL 01 | No evident peaks |
| SRR902003 | Spraguea lophii | No evident peaks |
| SRR902029 | Spraguea lophii | No evident peaks |
| SRR1596194 | Vairimorpha ceranae | Low coverage |
| SRR926322 | Vairimorpha ceranae | Low coverage |
| SRR926333 | Vairimorpha ceranae | Low coverage |

| | | |
|---|---|---|
| SRR540287 | Nematocida ausubeli | Low coverage |
| SRR501104 | Nematocida ausubeli | Low coverage |
| SRR501105 | Nematocida ausubeli | Low coverage |
| SRR1585328 | Nematocida ausubeli | Low coverage |
| SRR058693 | Nematocida ausubeli | Low coverage |
| SRR543736 | Nematocida ausubeli | Sample dominated by host |
| SRR489793 | Nematocida ausubeli | Sample dominated by host |
| SRR401825 | Nematocida ausubeli | Low coverage |
| SRR065290 | Nematocida parisii ERTm3 | Low coverage |
| SRR065291 | Nematocida parisii ERTm3 | No evident peaks |
| SRR065292 | Nematocida parisii ERTm3 | No evident peaks |
| SRR065288 | Nematocida parisii ERTm3 | No evident peaks |
| SRR014461 | Vairimorpha ceranae BRL01 | Low coverage |
| SRR040448 | Enterocytozoon bieneusi H348 | Low coverage |
| SRR18286426 | Ordospora colligata | Sample dominated by host |
| SRR18286429 | Ordospora colligata | Sample dominated by host |
| SRR18286428 | Ordospora colligata | Sample dominated by host |
| SRR18286427 | Ordospora colligata | Sample dominated by host |
| SRR18286425 | Ordospora colligata | Low coverage |
| SRR18286424 | Ordospora colligata | Low coverage |
| SRR18286423 | Ordospora colligata | Low coverage |

**Table S3: Ribosomal small subunit (SSU) sequences reconstructed from unassembled reads using PhyloFlash (with –emirge enabled).**

| Accession | Species | Reconstructed SSU sequence |
|---|---|---|
| SRR23214363 | *Nematocida ausubeli* | AGGTTGATTCTGCCTGACATAGACACTAGTCTCTCAGACT<br>AAGCCATGCAAAACGGGCAAACGGAAGGAAGCGTTGTAC<br>AGCTCAAAAGAACAGTTTCAACCGGCTTGGAGGAAAGGG<br>GCGGACATCCGAGGAAACCTTCGGCCAAGACGACGCACG<br>TCGGGGGGCATCCACGGAGAAGCAGGACGCCAGAGCAG<br>CGCAGGGCGCGGCCAAACGGGAGACTGTCCTATTAGCTA<br>GAAGGTAAGGTCAGGGCTTACCTTGGCGACCATGGGATA<br>CGAGGAATTGGGGTTTGGTTTCGGAGAGGGAGTGTGAGG<br>TGCGGCTCCTAGATCCAAGGATCGCAGCAGGCGCGAAAC<br>TTGCTCACTTCCCGGGGGGAGAAGCAGTGAGGAGACATG<br>CGGGGAAAGAGCGGGCAGCAAAGGAGCGCGCGAGGAGC<br>GATTGGAGGGCAAGACTGGTGCCAGCAGCCGCGGTAATA<br>CCAGCTTCAAAAGTGTCTGTGGTGTTTGTTGTGATTAAAAG<br>GTCCGCAGTCGGGGAGAGAGGCCGTGGGGAAAGCAGGC<br>GGCTCAAAGGGAGAGGCAAGCACGGAAAGAACGCGTCAA<br>GAGGGGCAGGCGGGGGCAGGAGAATTCAGCAGCCAGAG<br>GTGAAATTCGCAGACTTGCTGGGGGCGGACAGAGGCGAA<br>GGCGCCTGCCAAGGACGCTTTCATTGATCAGGGACGAAG<br>GCCGGAGGATCGAAGACGATTAGAGACCGTTGTAGTTCC<br>GGCGGTAAACGGTGCCGACACGGGTGTCTGCGGCGGCG<br>GCGCAGCGGGCGCCCGGAGAGAAATCGAGTGCGAGGGC<br>TTTGGGGAAAGTACAGTCGCAAGACGGAAACTTAAACGAA<br>ATTGACGGAAGGACACCACCAGGAGTGGAGCGTGCGGCT<br>TAATTTGACTCAACACGGGGCACTTTACCGCTGGAAGACG<br>CCAGAGGGATCGGCGCGAGATTTGGCAGAAAGTGGTGCA<br>TGGCCGCTCCTGGCCCGTGGGGTGACCTGTCAGGTTGAA<br>TCCGCTAACGGGCGCGATCCGCGCGCCTTGACTGCGCGG<br>CAGGAAGGCGGACGGCCGGCGCGCAGCCGGAGGAGGG<br>CGGGCGATAGCAGGTCCGTGATGCCCTTTGAAGCAGCGG<br>GCTGCACGCGCGCTACAGTGCGGGCAGGAGGGCCGCGC<br>CGAGAGGCGCGCGCAGAGCGGGCCGCAGGGGATGCCCG<br>CGAGGGAGCGCGGGCTGAACGCGGAATTCCCAGTACCC<br>GCGGGTCACCAGCCCGCGGAGACAGCGTCCCTGTCCTTT<br>GTACACACCGCCCGTCGCTATCTGAGATGGGCGCGCGCG<br>GCAAGCGCGGGGGCGCGCGAGCTGCCGCGCCTAGATTG<br>GATAAAAGTCGTAACAAGGTTTCCGTAGGAGAACCTGCGG<br>AAGGATC |
| SRR17622377 | Nematocida major | AGGTTGATTCTGCCTGACATAGACACTAGTCTCTCAGACT<br>AAGCCATGCAAAACGGGCAAACGGAAGGAAGCGTTGTAC<br>AGCTCAAAAGAACAGTTTCAACCGGCTTGGAGGAAAGGG<br>GCGGACATCCGAGGAAACCTTCGGCCAAGACGACGCACG<br>TCGGGGGGCATCCACGGAGAAGCAGGACGCCAGAGCAG<br>CGCAGGGCGCGGCCAAACGGGAGACTGTCCTATTAGCTA<br>GAAGGTAAGGTCAGGGCTTACCTTGGCGACCATGGGATA<br>CGAGGAATTGGGGTTTGGTTTCGGAGAGGGAGTGTGAGG<br>TGCGGCTCCTAGATCCAAGGATCGCAGCAGGCGCGAAAC |

| | | |
|---|---|---|
| | | TTGCTCACTTCCCGGGGGGAGAAGCAGTGAGGAGACATG<br>CGGGGAAAGAGCGGGCAGCAAAGGAGCGCGCGAGGAGC<br>GATTGGAGGGCAAGACTGGTGCCAGCAGCCGCGGTAATA<br>CCAGCTTCAAAAGTGTCTGTGGTGTTTGTTGTGATTAAAAG<br>GTCCGCAGTCGGGGAGAGAGGCCGTGGGGAAAGCAGGC<br>GGCTCAAAGGGAGAGGCAAGCACGGAAAGAACGCGTCAA<br>GAGGGGCAGGCGGGGGCAGGAGAATTCAGCAGCCAGAG<br>GTGAAATTCGCAGACTTGCTGGGGGCGGACAGAGGCGAA<br>GGCGCCTGCCAAGGACGCTTTCATTGATCAGGGACGAAG<br>GCCGGAGGATCGAAGACGATTAGAGACCGTTGTAGTTCC<br>GGCGATAAACGATGCCAACTCGGGGGCGTGCGGCCGGG<br>AGGGCGCGGGCGCCCGGAGAGAAATCGAGTTCGAAGGC<br>TTTAGGGAAAGTACAGTCGCAAGGCAGAAATTTAAACGGA<br>AATTGGCGGATTAATACACCAGGAGTGGAGCATGCGGCTT<br>AATTTGACTCAACACGGGGCAGCTTACCGCTGGAAGACG<br>CTGGGGGGATCGGGGCGAGATTTGGCAGAGAAGTGGTG<br>CATGGCCGCTCTTGGTCCGTGGGGTGACCTGTCAGGTTG<br>AATCCGCTAACGGGCGCGACGCGCGTGCGCCAAAAGAAA<br>GAGGGTGCAGACGGCCGGCGCGCAGCCGGAGGAGGGC<br>GGGCGATAGCAGGTCCGTGATGCCCTTTGAAGCAGCGGG<br>CTGCACGCGTGTTACAGTGTGCGCAGGAGGGGTGCGCCG<br>AGAGGCGCTGCAGCAAACGCGGGCGGGGGATGCCGGCG<br>AGGGAGCGCGGGCTGAACACGGAATTTCCAGTACCCGCG<br>GGTCACCAGCCCGCGGAGACAGCGTCCCTGTTTTTTGTAC<br>ACACCGCCCGTCGCTATCTGAGATGGGCGGCGCGGCAAG<br>CGCGGACACGCGCGAGCTGCTGCGCCTAGATTGGATAAA<br>AGTCGTAACAAGGTTTCCGTAGGAGAACCTGCGGAAGGAT<br>CACTTC |
| SRR16954902 | Hamiltosporidium magnivora | CACCAGGTTGATTCTGCCTGACGAGGATGCTTGTCTCTGG<br>GATTAAGCCATGCAAGTCTGGTGAAGCGAAAGTGGAACTG<br>CGAACGGCTCAGTAGAACGGTGATTATTTAATCGTGTAGG<br>AAGGATAACCGCGGGAAACTGTGGCTAATAACACGAGTAA<br>GACGCCGACCCATCAGTTTTATCGTACGGTAAGGGCGTAC<br>GATGGCTTTAACGGGTACGGGGGATCAGGGTTTGATTCC<br>GGAGAGGGAGCCTGAGAAACGGCTACCAGGTCTAAGGAC<br>AGCAGCAGGCGCGAAACTTGCCCAATTGGAGAGAGGCAG<br>TTATGAGACGTATATTTTGTAACCGGGGTAGAATACCGGT<br>GATTGACTGGAGGGCAAGTCTGGTGCCAGCAGCCGCGGT<br>AATTCCAGCTCCAGGAGTGTCATAGTGCATTGCTGCATTT<br>AAAAGGTCCGTAGTCAGTGTTGACAGGGATGCTAGAAGAC<br>ACTTTCTTCATGGAGTGTATTGGCATTCTAGGTTAGCATGT<br>AGGAGCGGAAGAGGGCGACTGTATTGCGTAGCGAGAGGT<br>GAAAGTTGACGACCTACGTAGGACAAACCGAAGTGAAAGC<br>TGTCGTCCAGTACGTTTCCGGTGATCAAGGACGTAAGCCG<br>GAGGAGCAAAGGTGATTAGAGACCCCTGTAGTTCCGGCC<br>GTAAACTATGCCAACTGGGTGTTACGTAGTGGTACCCACG<br>AGAAATCAAGTATATGGGCTATGGGGATAGTACGATCGCA<br>AGATTGAAACTTGAAGAAATTGACGGAAGGACACCACCAG<br>GAGTGGAGTGTGCGGCTTAATTTGACTCAACGCGGGACA<br>ACTTACCAGGGCCAGTTGTATGACGAATCTATGATGAGTA<br>CGACTAAGTGGTGCATGGCCGTTCACAACATGTGAGGTGA<br>CTTTTAGGTTTATTGCTGTAATGTGTGAGACCCTCAGCTAC<br>GGCGACTGGGGACTATAAGTTTCAGGAAGGAGGGGGCTA<br>TAACAGGTCTGTGATGCCCTTAGATGTTCTGGGCTGCACG<br>CGCACTACAATGTTATTTCTAGGTATATTGTAAAGATAAAG<br>AATAACTCGGTTGGGATTGCGTTCTGTAATGGACGCATGA<br>ACCAGGAATTCCTAGTAGTCGCGTGTCACTAACGGGCGAC<br>GACTGCGTCCCTGTTCTTTGTACACACCGCCCGTCGTTAT |

| | | |
|---|---|---|
| | | CTAAGATGGTGCTATGTTCGAAGAGGGTTTTCCTTCTGAG<br>GACGTAGGATTAGATTGGATACAAGTCGTAACAAGGTTGC<br>TGTAGGAGAACCTGCAGCAGGATCAGTGATAGTACTTTTA<br>TTGTGTAGTTTTTTCATATGAAACTCATACAAGGGATCACT<br>TGGCTCCGTATCCGAGGAAGGCCGGAATAGTTTCCGAGA<br>ACTAAGGCCGAACGGCATTTGTAAATGCCTGACTACCCCC<br>TGGACTTAAGCATATTATTAAGGGGAGGAACAAAAACTAA<br>CTAGGATTTCTGTAGTAGCGGCGAGCGAACAAGAAATAGC<br>CCCGAATGTAAGCCTCCGGGCATTGTTAAGATCGTCGTAA<br>CGAATTACCGGGACAGGTAAGCCATAGAGGGTGATAGCC<br>CCGTAGTGACGAGTAATGGGCAGAGTAGGGTTGCTTGGT<br>AATGCAGCCTGAAGAGGTGGTGGGTTCCATCTAAGGCTAA<br>ATATGAGCGGAGACCGATAGCGTAATAGTACAGCGATGGA<br>AAGGTGAAAAGCTAAAAAGTGCAAAGACGTGAAATTGTAT<br>TGAGTATCCTGATAACAGGACCCGTCTGAAACACGGACCA |
| SRR16954899 | Hamiltosporidium tvaerminnensis | AGGTTGATTCTGCCTGACGAGGATGCTTGTCTCTGGGATT<br>AAGCCATGCAAGTCTGGTGAAGCGAAAGTGGAACTGCGA<br>ACGGCTCAGTAGAACGGTGATTATTTAATCGTGTAGGAAG<br>GATAACCGCGGGAAACTGTGGCTAATAACACGAGTAAGAC<br>GCCGACCCATCAGTTTTATCGTACGGTAAGGGCGTACGAT<br>GGCTTTAACGGGTACGGGGGATCAGGGTTTGATTCCGGA<br>GAGGGAGCCTGAGAAACGGCTACCAGGTCTAAGGACAGC<br>AGCAGGCGCGAAACTTGCCCAATTGGAGAGAGGCAGTTA<br>TGAGACGTATATTTTGTAACCGGGGTAGAATACCGGTGAT<br>TGACTGGAGGGCAAGTCTGGTGCCAGCAGCCGCGGTAAT<br>TCCAGCTCCAGGAGTGTCATAGTGCATTGCTGCATTTAAA<br>AGGTCCGTAGTCAGTGTTGACAGGGATGCTAGAAGACACT<br>TTCTTCATGGAGTGTATTGGCATTCTAGGTTAGCATGTAGG<br>AGCGGAAGAGGGCGACTGTATTGCGTAGCGAGAGGTGAA<br>AGTTGACGACCTACGTAGGACAAACCGAAGCGAAAGCTGT<br>CGTCCAGTACGTTTCCGGTGATCAAGGACGTAAGCCGGA<br>GGAGCAAAGGTGATTAGAGACCCCTGTAGTTCCGGCCGT<br>AAACTATGCCAACTGGGTGTTACGTAGTGGTACCCACGAG<br>AAATCAAGTATATGGGCTATGGGGATAGTACGATCGCAAG<br>ATTGAAACTTGAAGAAATTGACGGAAGGACACCACCAGGA<br>GTGGAGTGTGCGGCTTAATTTGACTCAACGCGGGACAACT<br>TACCAGGGCCAGTTGTATGACGAATCTTTGATGAGTACGA<br>CTAAGTGGTGCATGGCCGTTCACAACATGTGAGGTGACTT<br>TTAGGTTTATTGCTGTAATGTGTGAGACCCTCAGCTACGG<br>CGACTGGGGACTATAAGTTTCAGGAAGGAGGGGGCTATA<br>ACAGGTCTGTGATGCCCTTAGATGTTCTGGGCTGCACGCG<br>CACTACAATGTTATTTCTAGGTATATTGTAAAGATAAAGGA<br>TAACTCGGTTGGGATTGCGTTCTGTAATGGACGCATGAAC<br>CAGGAATTCCTAGTAGTCGCGTGTCACTAACGGGCGACG<br>ACTGCGTCCCTGTTCTTTGTACACACCGCCCGTCGTTATC<br>TAAGATGGTGCTATGTTCGAAGAGGGTTTTCCTTCTGAGG<br>ACGTAGGATTAGATTGGATACAAGTCGTAACAAGGTTGCT<br>GTAGGAGAACCTGCAGCAGGATCA |
| SRR8476225 | Astathelohania contejeani | GGTTGATTCTGCCTGACGTGGAAGCTATTCTTTAAGATTAA<br>GCCATGCATGTGTAGAATGAAGTGAAGCCATTAGGTGGAA<br>CAGCGAAAAGCTCAGTAATACAATCATTATTTGGTCTAC<br>AAGATATAGAATAACCTTGATAAATTAAGGCTAAAGCTATT<br>GTAGAATAAGAGATTGACCTATCAGCTAGTATGTAGGGTA<br>AGGGCCTACGTAGGCGATGACGGGTAACGGGGAATTAGG<br>GTTCTATTCCGGAGAAGGAGGCTGAGAGATGGCTACTAG<br>GTCTAAGGAGAGCAGCAGGCGCGAAACTTACCCAATGCT<br>ATTTAGTAGTGTAGAGTGTAGTGAATATTCTAATGGAGAGG |

| | | |
|---|---|---|
| | | CTAGTAAAGCAACCATTGTAATTCCAGCAAATCTATTACGA<br>GTATTGCTGCAATTAAAAAGTTCGTAGTTGATTATTGTAAT<br>AATCTTGTGATAAGTATTAAATTATGATTAATGAAAGCCAT<br>GGAAGGAAATAGGATTAAATAGGGAGGGGTGAAATCTGTA<br>GATCTATTTAGGACTAACTGAAGCGAAAGCGATTTTCTATG<br>TGGTATTTGACAATCAAGAACGAAAGCCGGAGTATCGAAG<br>ACGATTAGAGACCGTCGTAGTTCCGCGCAGTAAACTATGT<br>TATATTATTTGTAGTATATATGAGTATATATTATGGATATAT<br>AGAAATTAAGATATTATGAACTTTGGGGATAGTACGAACGC<br>AAGTTTTAAACTTAAATGAAATTGACGGAAGGACACACCAG<br>GAGTGGAGTGTGCGGTTTAATTTGACTCAACGCGGGACAA<br>CTTACCATTTTTAGAAGTGAATATGAATGATATTTATCATGA<br>TTTTACTATGAGTGGTGCATGGCCGTTAACAATACGTGAT<br>GTGAATTTTGAATTTGAATGAGTTATATATGAAAGTATATAA<br>TTCTGTATTAGTGTTAAATCCACCAATGTGTGAGACCCATA<br>TACTAATTTTGAGTTAGTAGACAGATGATGTAAATCATAGG<br>AAGGGATGGGCGATAACAGGTCAGTGATGCCCTTAAATGA<br>AATGGGCGACACGCGCACTACAATAGAATGATATATTATT<br>CTAAGGAGTTGGGATTATTAATATGTAAATTTAATATGAAC<br>AAGGAATTCCTAGTAATATTTTTTTTGTGATGTTAACAACGA<br>GATATGAATATGTCCCTGTCCTTTGTACACACCGCCCGTC<br>GTTATCTCAGATGGATATTAGGGTGAAATATTAATATAATA<br>GAGAACTCTAATAACTAAATAAGGTACAAGTCGTAACAAG<br>GTTGACCTAAGTGAACTTGGGGCAGGATCA |
| ERR3154977 | Tubulinosema<br>ratisbonensis | GATTGATTCTGTCTGTTATATGTGCTAGTGTCGAAGATTTA<br>GCCATGCATGCTTTACGAATTCACAAGAAGAAGTGGCGGA<br>CAGCTCAGTAATACAGTTATAATATACTCTCTACTAAAGGA<br>TAACCGCGGTAAGCTGCGGGTAAAACTTCAAGCTATGTGT<br>TTCATGAAAGAATTTATAGTTCAGAGTGTTAGTTTCTGGCC<br>TATTAGTTAGTAGGTAATGTAAGGGATTACCTAGACTATTA<br>TGGGTAACGGGAGATGAATGTCTGATACCGGAGAGGAAG<br>CCTTAGAAACCGCTTTCACGTCCAAGGATGGCAGCAGGC<br>GCGAAACTTACCCAATCGTTTTTTAAGACGGAGGTAGTTAT<br>GACACATGTGAATTATTCACGAGGAAGATCAATAAATCAGA<br>TTTTTTTTACTGGAGGGCAAGTCTGGTGCCAGCAGCCGCG<br>GTAATTCCAGCTCCAGTAGTGCATATACATGCTGTAGTTAG<br>AAAGTTTGTAGCCGTATAAGCTTGAATCAGAGGAAAGGAG<br>GACTCTTAAGGTAAACTTTAACTCTGGTCAATGCTTATATT<br>CGAAGCGGATGGAGGCACTTGTATTCAATAGCGAGAGGT<br>AAAATTTGATGACCTATTGAGGACAATCAGTAGCGAAAGC<br>GAGTGTCTAGTACGCGTTTGAGGGTCAAGAACGTTAGCCG<br>GAGGATCGAAGATGATTAGATACCGTTGTAGTTCCGGCCG<br>TAAACCATGCCTACTTGCCTTGTACTTAGTATGAAGCATAG<br>AGAAATTAAGAGTTTTTGGGCTCTAGGGATAGTAATCCGG<br>CAACGGACAAACTTAAAGAAATTGGCGGAAGGACACCACA<br>AGGAGTGGATTGTGCGGCTTAACTTGACTCAACACGGGAA<br>ATCTTACCAGGGTCTGTATATGTGAGACTGTCCCATTTATG<br>GTGGTACACGATAATATACTGAGTGGTGCATGGCCGTTTT<br>CAACACGTGGGGTGACCTGTCAGGTTTATTCCGGTAACGT<br>GTGAGGTCACAGTATTAATAAGTATTTTAATAGACAGCTAA<br>TGGTAAATTAGAGGAAGTGTGACGATAACAGGTCAGTGAT<br>GCCCTTAGATACCCTGGGCTGCACGCGCAATACATTGAGT<br>AAGTCATTTACAACAGTAATTTTGAAACTTACTCGCTACTG<br>GGATCATCCTTTGTAATTGGGATGTGAACGTGGAATTCCT<br>AGTAATCGTAGCTCACTAAGTTACGATGAATGTGTCCCTGT<br>TCTTTGCACACACCGCCCGTCGCTATCTAAGATGGATGTT<br>GATATGAAATTGCTGCTTGAATGTGAGGTACTTGAGTATTA |

| | | |
|---|---|---|
| | | ACAACTAGATAAGATATAAGTCGTAACAAGGCTGCTATAGA<br>AGAATCTGTGGCAGGATCA |
| SRR489790 | Annaliia algerae | CACCAGATTGATTCTGTCTGGTATGTGTGCTAGCGTCAAA<br>GATTTAGCCATGCATGCTTTTCGAACCCTCGTGGGGAGAG<br>GCGGATAGCTCAGTAATACAGTTATAACATAAGCTGCGTG<br>TGTGGATAACCTTGTTAAGATAAGGCTAAGACTTAATAAGT<br>CGCACTTTTGTGAAGAAACGCGACTTGTGCAGCATTGGTT<br>TCTGACCTATCAGTTAGTATGTTCTGTAAGGGAGAACATAG<br>ACTATGACGGGTAACGGGGGATGCACGTCTGATACCGGA<br>GAGGAAGCCTTAGAGACAGCTTTCACGTCCAAGGATGGC<br>AGCAGGCGCGAAACTTACCCAATTGTTTTTTGTGACAGAG<br>GTAGTTATGACGTATTCTTTAGAGAGAGACCTTGTTAGACA<br>TGGTCCATAGCGACTGGAGGGCAAGTCTGGTGCCAGCAG<br>CCGCGGTAATTCCAGCTCCAGTAGTGCATATACATGCTGT<br>AGTTAGAAAGTTTGTAGCCTATTTATGGATTGTTTTAGACA<br>AAAGGACGACTCAAATTGACCTTTCATTTGACTAATGCATG<br>AATGTAGAAGCGATTGAAGGCGATTGTATTCACCAGCCAG<br>AGGTAAAATTTGATGACCTGGTGAGGACACACGGAGGCG<br>AAAGCGATTGCCTAGAGCGTATTCAGTGGTCAAGAACGTA<br>AGCCGGAGGATCAAAGATGATTAGATACCGTTGTAGTTCC<br>GGCCGTAAATTATGCCAACTTGTGCTTCTGCTTCTGCGGA<br>GGCGCATAGAGAAATCAAGAGTTTATGGGCTCTAGGGATA<br>GTAATCCGGCAACGGACAAACTTAAAGAAATTGGCGGAAG<br>GACACCACAAGGAGTGGATTATGCGGCTTAATTTGACTCA<br>ACGCGGGACAACTCACCAGAGCCTATGTGCAGGAGACAG<br>TGAGCTTTGAGAGCGGACTGGATAGTACTTTGAGTGGTGC<br>ATGGCCGTTTGCAACACGTGAGGTGACTTGTCAGGTTTAC<br>TCCGGTAACGTGTGATGTGCTGTATGCAAGTATTTTTGTGA<br>GACTGCAGGCGGTAAGCCTGATGAAGCGGCGCTATAACA<br>GGTCAGTGATGCCCTTGGATGTTCTGGGCTGCACGCGTA<br>ATACAGTGGGAGCTGTAGATATGTATAGGTGGAAAAGTTC<br>CCGAGACTGGGATCATGCTTTGTAAGAAGGATGTGAACGT<br>GGAATTCCTAGTAATCGCTGCTCACTAAGTAGCGATGAAT<br>GAGTCCCTGTTCTTTGCACACACCGCCCGTCGCTATCTGA<br>GATGGATGTTTTTATGAAGATGCTGCTGTTAGAGGCATTTG<br>AGTAAGGACGACTAGATTAGATATAAGTCGTAACAAGGCA<br>GCGGTAGAAGAATCTGCCGCTAGATCATAAT |
| SRR8495097 | Cucumispora<br>dikerogammari | CAGGTTGATTCTGCCTGACGTGGACGCTAGTCTCATAGAT<br>TTAGCCATGCATGTGTAAGCGAACAAGAGGAAGCTGCGG<br>ACTGCTCAGTAACAGACATATAATTTAATCTTTACAGAAAC<br>GAGCGGAATAAACTCAGGAAACAGAGTGCAATACGTAAAA<br>GACGAATTTTTTATTATAAGAAATACGTTTTTAGCTTGAACA<br>AAGCGGTAAAGAATAAGTTGTCAGCCTATCAGTTAGTAAG<br>TAGGGTAAGGGCCTATTTAGACGAAGACGGGTACGGGGA<br>ATTAGAGTTTGATTCCGGAGAGGGAGCCTGAGAAATAGCT<br>ACCAGGTCCAAGGACGGCAGCAGGCGCGAAAATTACCGA<br>AGCTCGAATAGAGGCGGTAGTAATGAGACGTATTAATATA<br>AAACAAGGGTAAAAAACTTGTTAGTAACTGGAGGTCAAGT<br>CTGGTGCCAGCATCCGCGGTAATACCAGCTCCAGGGGTG<br>TCTATGATGATTGCTGCGATTAAAAGGTCCGTAGTCGAATT<br>TATATAATTGTTTGTAATATGCTAGATAAAATAACAGAAAGA<br>ACAATTACTTTAAATGAAAGGAATAGTAAGGGGCTGATTAA<br>TTGAGCAACGAGAGGTGAAATTTGATGACTTGCTTAGGAG<br>AAACAGAGGCGAAAGCGTCAGTCAAGTATAAATCCTATGA<br>TCAAGGACGTAGGCTAGAGTATCGAACACGATTAGATACC<br>GTAGTAGTTCTAGCAGTAAACTATGCCTACACTATCGAATA<br>AAAGTTTGGTAGAAGAGAAATCTTAAGTAGGGCTTTGGGG |

| | | |
|---|---|---|
| | | AGAGTACACGCGCAAGCGATAAATTTAAAGGAAATTGACG<br>GAAGAACACCACAAGGAGTGGAGTGTGCGGCTTAATTTGA<br>CTCAACGCGGGACAGCTTACCATACCCGAGGACTATAAGA<br>GTGAATACACGATAAGTCTAAAAGTGGTGCATGGCCGTTA<br>TCGACGAGTGAAGTGATTTTATGGTTAAATCCGACAAGTT<br>GTGAGACCCTTATTTAAATACAGGTATTGTTAAAATACAGG<br>AAGGAAAGGACAAGAACAGGTCAGTGATGCCCTTAGATG<br>GTATGGGCTGCACGCGCACTACAATGGTTATAATAATAAA<br>GATAATTAAAGTATAAATATAATCAAGAGGAATTGAGAACT<br>GAAAAGTTCCCATGAACGAGGAATTGCTAGTAATCGTAGG<br>CTCAGTAAGATACGATGAATATGTCCCTGTTCTTTGTACAC<br>ACCGCCCGTCGTTATCGAAGATGGAGTTTTACCCGAACAA<br>GCTTAAGCGAGTGAGTGTATGATTCTAGATCTGATACAAG<br>TCGTAACAAGGCAGCCGTAGGAGAACCTGCTGCTGGATC<br>AACT |
| SRR926312 | Pseudoloma neurophilia | CAGGTTGATTCTGCCTGACGTGGATGCTAGTTTCATAGAT<br>TAAGCCATGCATGTGTAAGCGAAGCGTAAGTGGAGCGGC<br>GTACGGCTCAGTAACGGGCTACTATTTGATCTCCCTGGGC<br>GGATATCCTCTGTAAACGGAGGGCAAAACGCAAGACAAG<br>CAGCAATTTTGTTGGCTGTCTAACGAAAGTGGGGGAGAGT<br>AAGGAGCCAGCCCATCAGCTAGTAAGTAGGGTAAGGGCC<br>TACTTAGGCAAATACGGGTACGGGGAATTATCGTTTGATT<br>CCGGAGAGGGAGCCTGAGAGATGGCTACCGGGTCCAAG<br>GACAACAGCAGGCGCGAAAATTACCGGAGCCTGAAGTCA<br>GGGCGGTAGTAAGGAGACGTGAGAACAATGTGGGGGTAA<br>AAAACGCACTGGAAACAGGAGGAAAAGACTGGTGCCAGC<br>ACCCGCGGTAATACCAGCTCCTGGAGTGTCTATGGTGATT<br>GCTGCAGTTAAAGCGTTCGTAGTCGGAGCGGAAAGAATG<br>GCGTGACAGACAGCTGCTCAAAGGGTGGTGTGCGCCGTG<br>ATCCCGCGGAATGAGGAGAGTTTTGGGACCAGGCTATTAA<br>ACGGCAAGCGGTGAAATGTGGTGACCCGTTTAGGAGCAA<br>CAGAGGCGAAAGCGCTGGTCAGGGGCTATTCCGATGATA<br>AAGGACGTAGGCTAGAGGATCGAAGACGATTAGAGACCG<br>TTGTAGTTCTAGCAGTAAACGATGCCGATGTTGTGGTGCA<br>GAGTGCAACGCAAGAGAGAAATCTAGTAGGGCCCTGGGG<br>AGAGTACGCGCGCAAGCGAGAAATTTAAAGGAAATTGACG<br>GATGAACACCTCAAATAGTGGAGTGTGCGGCTTAATTTGA<br>CTCAACGCGGGACATCTTACCGGGCCCGACGACCGGACG<br>AGCGTGACACGCGATAGGTCGGAGAGTGGTGCATGGCCG<br>TTAACGACGAGTGGAGTGATCTTTGGGTTAAGTCCGTAAA<br>TTAGTGAGACCCCAGCGGAAGGACAGGTGCGCAACGCAC<br>AGGAAGGATGGGTCAAGGACAGGTCAGTGATGCCCTTAG<br>ATGGCCCGGGCTGCACGCGCACTACAGTGGTCGCCGAAA<br>TTTGGAGAGAGAGGGAAAGGCGATCGAGAGGGAATGAGC<br>TTTGGAAGAGGCTCAGGAACGCGGAATTGTTAGTAATCGC<br>GGGCTCACTAAGACGCGATGAATGCGACCCTGTTCATTGT<br>ACACACCGCCCGTCGTTATCGAAGACGATGCTAGGCGCG<br>AGCAAGGTTTATTTGGCTGAGCGAGCGCAGGGTATTGAAT<br>CTGATGTAAGTCGTAACAAGGTAGCTGTAGGAGAACCTGT<br>AGCTGGATCA |
| SRR926341 | Agmasoma penaei | CACCAGGTTGATTCTGCCTGACGTAGAGGCTAGCCTCAG<br>GGACTAAGCCATGCATTGTTAGTGAAGTTTTTAATGAAACG<br>ACGGACGGCTCAGTAATACTACTTTTAACTAACCTTTTGTA<br>CTAATAATTAAGGGAAACTGTAATTAAAAATCATGAGGATG<br>TGAGGTAGACCTATTAGCTAGTTGGTTGTGTAAAGGACTA<br>CCAAGGCTATAATGGGTAACGGAGATTTAGTGATCGAAAC<br>CGGAGATGGAAGCTGAGAAACGGTTCCAATGTCCAAGGA |

| | | |
|---|---|---|
| | | TAGCAGCAGGCGCGAAAATTGCACACTCTTTAATGGGGAT<br>GCAGTTATGAGGTATGACAGAAAGGGTTATCAATAAATAA<br>GATGACGTAAAGCTATTAGAGGGAAAGTTTGGTGCCAGCA<br>GCCGCGGTAATACCAACTCTAAGAGTCTCTATGCGAGTTG<br>CTGCAGTTAAAAAGTCCGTAGTCTTTACGTAATTAAAAATG<br>AATGATCAAGTTTCATATTTTTACGTTTATATATGAGACGGA<br>TTGGGAGCATAGTATAACTGGGTTAAGAATGAAATCTCACT<br>ACCCTAGTTGGACTATCAGAAGCGAAAGCGATGCTCTAAT<br>ACGTACTTTTAGATAAAGGACGAAGGCTAGAGTAGCGAAA<br>GGGATTAGATACCCCTGTAGTTCTAGCAGTAAACTATGCC<br>GACAGAATGTTAGATATATTTCTAGTGTTCAAGGGAAACCT<br>TAAGTGATCGGGCTCTGGGGAGAGTATGCTCGCAAGTGT<br>GAAAATTAAACGAAATTGACGGAGTTACACCACAAGGAGT<br>GGATTGTGCGGCTTAATTTGACTCAACGCGAGGAATTTTA<br>CCAGGGCTGAATATATTTGAGATTGATTACATGAAATATAT<br>TTGAGTGGTGCATGGTCGTTGTAAACTCATGGATTGATCTT<br>AAGTTCAACTGCTAAAATGGGTGAGACTTTCATAAACAGCT<br>ATCTAACAGGTAGAGGAAGGGGAAGGCGATAACAGATCC<br>GTGATGCCCTCAGATGTCCTGGGCTGCACGCGCAATACAT<br>TATGTATATTTCTTATAAATAGATACTACATATTGGGGAATT<br>GACTTTTGTAAATAAGTCATGAACTTGGAATTCCTAGTAAT<br>AATGATTCATCAAGTCATTGTGAATGTGTCCCTGTAGCTTG<br>TACACACCGCCCGTCACTGTCTCAGATGGTTGATGAGATG<br>AAGAGCTTCGGTTCTGAATCTTAACAACTAAATAAGACATA<br>AGTCGTAACAAGGTATCG |
| SRR23214350 | Pancytospora philotis | CGGCGACCGAACGGCGAACGGCTCAGTAATACTGCGACG<br>ATCTGCCCCGCGCGGCCGATAACCGCGGGAAACTGCGGC<br>TAAGAGCGCGGGTTGAGACGCAGCCCTATCAGCTCGTTG<br>GCGGTGTAACGGACCCCCAAGGCGGCGACGGGTGACGG<br>GGGGTCGGGGCCCGACTCCGGAGAGGGAGCCTGAGAGA<br>TGGCTCCCACGTCCAAGGACGGCAGCAGGCGCGGAAATT<br>GCCCACTCCCGCGCGGGGAGGCAGTTACGAGACGTGGC<br>GGAAGGGCGCCCCGCAACGCCGGGCGCGAAGGCGATTG<br>GAGGGCAAGCTCGGTGCCAGCAGCCGCGGTAATACCGAC<br>TCCAAGAGTGTCTATGGTGGATGCTGCAGTTAAAACGTCC<br>GTAGTCGGCACGGCAACGAAGACGCGCGGCTCGAACGC<br>GCGCGCGTTGCCGGACGCGGCGGAGCGGGCAGGGGGC<br>GCGGTATGCGCGGGCGAGAGATGAAACGCCAGGACCCC<br>GCGCGGACCGACGGCGGCGAAGGCGGCGCCCCGGGAC<br>GCGTCTGCGGATCAAGGACGAAGGCCGGAGGATCGAAAG<br>TGATTAGAGACCGCTGTAGTTCCGGCAGTAAACGATGCCG<br>ACAGCGCGGCGCGCGTGCGCCGCGCGGGGGAAACCTTA<br>GTGTGCGGGCTCTGGGGATAGTATGCTCGCAAGCGTGAA<br>AATTAAACGAAATTGACGGAGCTACACCACAAGGAGTGGA<br>TTGTGCGGCTTAATTTGACTCAACGCGGGGCAGCTTACCA<br>GGGCCGCGCGCGCCCGAGACCGCGCGCGCGAGGCGCG<br>CAGGAGTGGTGCATGGTCGTTGCAAGCCGATGGCGCGAG<br>CTTAGGCTTAAGTGCCGGAATGGGCGAGATCCGCGCGAC<br>GGGTGCGCGCCGCACAGGCGGGAGCGGGCGATAACAGA<br>TCAGTGATGCCCTCAGATGCCCTGGGCCGCACGCGCAAT<br>ACACTCCCCGCGCGCGGCGCAGACGGGTCGCGGGGGGC<br>GGGACCGGCGCCTGCAAGGGCGCCGCGAACGAGGAATT<br>CCTAGTAACGGCGGCTCACCAGGCCGCGGTGAATGTGTC<br>CCTGTAGCTTGTACACACCGCCCGTCACTATCTCAGACGG<br>CCGCCGCGGCGAAGGGAGCG |

| | | |
|---|---|---|
| SRR065293 | Vittaforma corneae | ACCAGGTTGATTCTGCCTGACGTAGATGCTAGTCTCTAAG<br>ATTAAGCCATGCATGTTTCCGCAATCAGGGACGAATAGCT<br>CAGTAAAACTGCGATGATTTAGTCTGGCTGTGTAGATAACT<br>ACGTGAAAATGTAGCTAAGGGAAGGCAGAATAAGACGCA<br>GGACTATCAGTTAGTTGGTAGTGTAATGGACTACCAAGAC<br>AGTGACGGTTGACGGGAAATTAGGGTTTTGTACCGGAGA<br>GGGAGCCTGAGAGATTGCTCCCACGTCCAAGGACGGCAG<br>CAGGCGCGAAAATTGCCCACTCTTTGCAGGAGGCAGTTAT<br>GAGACGTGAAGATGAGTATCTTGTAAAGAGGGATAGGAGA<br>ATTGGAGGGCAAGTTTGGTGCCAGCAGCCGCGGTAATAC<br>CGACTCCAAGAGTGTGTATGAGAGATGCTGCAGTTAAAAA<br>GTCCGTAGTCATAGAAGGGCAAAGAGAGATGCGAGGTCT<br>CACAGTGCGATGATGGAGGAGCCGATGGGGAACATAGTA<br>TACCAGGGCGAGAGATGAAATGCCAAGACCCCTGGTGGA<br>CTGAGCGAGGCGAAGGCGATGTTCTTGTAGGCATTCGGT<br>GATCAAGGACGAAGGCTGGAGTATCGAAAGTGATTAGATA<br>CCGCAGTAGTTCCAGCAGTAAAAGATGCCGACATGCTCAT<br>TGGACACAGTGGGCAGGGAGAAATCTTAAGAGTTCGGGC<br>TCTGGGGATAGTATGCTCGCAAGGGTGAAAATTAAAGAAA<br>TTGACGGAGCTACACCACAAGGAGTGGATTGTGCGGCTTA<br>ATTTGACTCAACGCGAGGAAACTTACCAGGGCCAAGTATT<br>GTGTAGAAACGAGCAATACAGGAGTGGTGCATGGTCGTT<br>GGAAATTGATGGGATGACTTTGACCTTAAATGGTTGAATG<br>AGTGAGATCTTTTGGACATGTTCCGCACGGAACAGGAAGG<br>AAAAGGCTATAACAGATCCGAGATGCCCTCAGATGCCCTG<br>GGCTGCACGCGCAATACAATAGCAGGTAGAGAGAGAGAC<br>AGGAAGGTGCTCAGATGGACTATGTGCTGTAAGGCACATA<br>CGAAAGAGGAATTCCTAGTAAGTGTGTATCAACAATGGAT<br>ATTGAATAAGTCCCTGTAGCTTGTACACACCGCCCGTCAC<br>TATCTCAGATGTTTTTCAGGATGAAGAGTCCAGGCTCTGA<br>ATAATGAAAAGTAGATAAGATGTAAGTCGTAACAAGGTTGC<br>GGTCGGTGAACCAGCCGCAGGATCATTG |
| SRR17317295 | Vairimorpha ceranae | CACCAGGTTGATTCTGCCTGACGTAGACGCTATTCCCTAA<br>GATTAACCCATGCATGTTTTTGACATTTGAAAAATGGACTG<br>CTCAGTAATACTCACTTTATTTTATGTAAATTTTTAATTAACT<br>ACGTTAAAGTGTAGATAAGATGTTTACAGTAAGAGTGAGA<br>CCTATCAGCTAGTTGTTAAGGTAATGGCTTAACAAGGCTG<br>TGACGGGTAACGGTATTACTTTGTAATATTCCGGAGAAGG<br>AGCCTGAGAGACGGCTACTAAGTCTAAGGATTGCAGCAG<br>GGGCGAAACTTGACCTATGGATTTTATCTGAGGCAGTTAT<br>GGGAAGTAATATTATATTGTTTCATATTTTAAAAGTATATGA<br>GGTGATTAATTGGAGGGCAAATCAAGTGCCAGCAGCCGC<br>GGTAATACTTGTTCCAAGAGTGTGTATGATGATTGATGCA<br>GTTAAAAAGTCCGTAGTTTATTTTTAAGAAGCAATATGAGG<br>TGTACTGTATAGTTGGGAGAAAGATGAAATGTGACGACCC<br>TGACTGGACGAACAGAAGCGAAAGCTGTACACTTGTATGT<br>ATTTTTTGAACAAGGACGTAAGCTGGAGGAGCGAAGATGA<br>TTAGATACCATTGTAGTTCCAGCAGTAAACTATGCCGACG<br>ATGTGATATGTATTAATTTGTATTACATAATAGAAATTTGAG<br>TTTTTTGGCTCTGGGGATAGTATGATCGCAAGATTGAAAAT<br>TAAAGAAATTGACGGAAGAATACCACAAGGAGTGGATTGT<br>GCGGCTTAATTT<br>GACTCAACGCGAGGTAACTTACCAATATTTTATTATTTTGA<br>GAGAACGGTTTTTTGTTTGAGAATGATAATAGTGGTGCATG<br>GCCGTTTTCAATGGATGCTGTGATTAATTTCAACAAGACGT<br>GAGACCCTTATTTTTTATTAAAGACAGACACAATCAGTGTA<br>GGAAGGAAAGGATTAAAACAGGTCCGTTATGCCCTCTGAC<br>ATTTTGGGCTGCACGCGCAATACAATAGATATATAATCTTT |

| | | |
|---|---|---|
| | | ATGGGATAATATTTTGTAAGAGATATTTGAACTTGGAATTG<br>CTAGTAAATTTTATTAAATAAGTAGAATTGAATGTGTCCCT<br>GTTCTTTGTACACACCGCCCGTCGCTATCTAAGATGATATA<br>TGTTGTGAAATTAGTGAAAACTACTTAAACAATATGTATTA<br>GATCTGATATAAGTCGTAACATGGTTGCTGTTGGAGAACC<br>ATTAGCAGGATCATAATGATTTTTAAAATTATTTATTTCATA<br>TTATTATTTTTATTTTGCCCACACATGGGATCAATAGGATA<br>CCATAACGATGAAGGTCGTAATAGAATACGAAAGTATTTTA<br>ATATTTACCGAATTAATATTTAATAATATTGATTACCCTT |
| SRR24007515 | Encephalitozoon intestinalis | CATCAGGTTGATTCTGCCTGACGTGGATGCTATTCTCTGG<br>GACTAAGCCATGCATGTTGATGAACCTTGTGGGGGATTGA<br>CGGACGGCTCAGTGATAGTACGATGATTTGGTTGGCGGG<br>AGAGCTGTAACTGCGGGAAACTGCAGGTAGGGGGCTAGG<br>AGTGTTTTTGACACGAGCCAAGTAAGTTGTAGGCCTATCA<br>GCTGGTAGTTAGGGTAATGGCCTAACTAGGCGGAGACGG<br>GAGACGGGGGATCGGGGTTTGATTCCGGAGAGGGAGCCT<br>GAGAGATGGCTACTACGTCCAAGGATGGCAGCAGGCGCG<br>AAACTTGCCTAATCCTTTGGGGAGGCGGTTATGAGAAGTG<br>AGTTTTTTTCGAGTGTAAAGGAGTCGAGATTGATTGGAGG<br>GCAAGTCGGGTGCCAGCAGCCGCGGTAATACCTGCTCCA<br>ATAGTGTCTATGGTGAATGCTGCAGTTAAAAAGTCCGTAG<br>TCTTTTGTATGTCTTTGTTTGGGGGATTATGTCCTGATGTG<br>GATGTAAGAGGTTTGGCAGAGGACGAGGGGCACCGGATA<br>GTTGGGCGAGGGGTGAAATACGAAGACCCTGACTGGACG<br>GACAGAAGCGAAGGCTGTGCTCTTGGACTTATGTGACGAT<br>GAAGGACGAAGGCTAGAGGATCGAAATCGATTAGATACC<br>GTTTTAGTTCTAGCAGTAAACGATGCCGACTGGACGGGAC<br>TATATAGTGTTGTGCATGAGAAATCTTGAGTATGTGGGTTC<br>TGGGGATAGTATGCTCGCAAGAGTGAAACTTGAAGAGATT<br>GACGGAAGGACACCACAAGGAGTGGAGTGTGCGGCTTAA<br>TTTGACTCAACGCGGGGCAACTTACCGGTTCTGAAGCGG<br>GCAGGAGAACGAGGACGGGATGCGCGCGGCGGTGGTGC<br>ATGGCCGTTTGAAATGGATGGCGTGAGCTTTGGATTAAGT<br>TGCGTAAGATGTGAGACCCTTTGACAGTGCTCTTTGGGGC<br>AAGGGAGGAATGGAACAGAACAGGTCCGTTATGCCCTGA<br>GATGAAGCGGGCGGCACGCGCACTACGATAGATGGCGAG<br>GGAGCCTGCTGTGAGGGATGAAGCTGTGTAATGGGCTTC<br>TGAACGTGGAATTCCTAGTAATAACGATTGAACAAGTTGTT<br>TTGAATGGGTCCCTGTCCTTTGTACACACCGCCCGTCGCT<br>ATCTAAGATGACGCAGTGGACGAAGATTGGAAGGTCTGAG<br>TCCTTCGTGTTAGATAAGATATAAGTCGTAACATGGCTGCT<br>GTTGGAGAACCAGCAGCAGGATCAGTATTTG |
| SRR23560257 | Encephalitozoon hellem | CATCAGGTTGATTCTGCCTGACGTGGATGCTATTCTCTGG<br>GGCTAAGCCATGCATGTTTATGAAGCCTTTATGGGGGATT<br>GACGGACGGCTCAGTGATAGTACGATGATTTGATTGGGAG<br>CCTGGATGTAACTGTGGGAAACTGCAGGTAAGTTCTGGG<br>GGTGGTAGTTTGTAGCTACTGCGTACCGAGTAAGTTGTAG<br>GCCTATCAGCTGGTAGTTAGGGTAATGGCCTAACTAGGCG<br>G<br>AGACGGGAGACGGGGGATCAGGGTTTGATTCCGGAGAGG<br>GAGCCTGAGAGATGGCTACTACGTCCAAGGATGGCAGCA<br>GGCGCGAAACTTGCCTAATCCTTATTGGGGAGGCGGTTAT<br>GAGAAGTAAGATGTTTAGCAAGTATAAATTTGTTGTGATTT<br>ACTGGAGGGCAAGTCGGGTGCCAGCAGCCGCGGTAATAC<br>CTGCTCCAGTAGTGTCTATGGTGAATGCTGCAGTTAAAAT<br>GT<br>CCGTAGTTGTTTGTATGTCTTTTGAGTGATGTTTATGGTTT |

| | | |
|---|---|---|
| | | TTAGTGGATGTAGTTTTATTGTAGCAGAGGACGAGGGGCA<br>CTGGATAGTTGGGCGAGGGGTGAAATACGAAGACCCTGA<br>CTGGACGAAGAGAAGCGAAGGCTGTGTTCTTGGACTTTTG<br>TGGTGATGAAGGACGAAGGCTAGAGGATCGAAATCGATTA<br>GATACCGTTTTAGTTCTAGCAGTAAACGATGCCGACTGGA<br>CGGGACTGTTTTAGTGTTGTCCGAGAGAAATCTTAAGTAT<br>GTGGGTTCTGGGGATAGTATGCTCGCAAGAGTGAAACTTG<br>AAGAGATTGACGGAAGGACACCACAAGGAGTGGAGTGTG<br>C<br>GGCTTAATTTGACTCAACGCGGGGCAACTTACCGGTTCTG<br>AAGTGAGTGTGAGAGTGTTTTTACATGATGCTTACGGCGG<br>TGGTGCATGGCCGTTTTAAATGGATGGCGTGAGCTTTGGA<br>TTAAGTTACGTAAGATGTGAGACCCTTTTTGACTGTGCTCT<br>ATGGGGCAAGGGAGGAATGGAACAGAACAGGTCCGTTAT<br>GCCCTGAGATGAAGCGGGCGGCACGCGCACTACGATAGA<br>TGCCTATGTGGGCTACTGTGAGGGATGAAGCTGTGTAATG<br>GGCTTCTGAACGTGGAATTCCTAGTAAGAATGATTGAACA<br>AGTTATTTTGAATGTGTCCCTGTCCTTTGTACACACCGCCC<br>GTCGCTATCTAAGATGACGCAGTGGACGAAGATTGAGAG<br>GTCTGAGTCTTTCGTGTTAGATAAGATATAAGTCGTAACAT<br>GGCTGCTGTTGGAGAACCAGCAGCAGGATCAGTA |
| SRR17858635 | Encephalitozoon cuniculi | CCAGGTTGATTCTGCCTGACGTGGATGCTATTCTCTGGGG<br>CTAAGCCATGCATGCTTGTGAACTCTTTGTGGGGGATTAG<br>CGGACGGCTCAGTGATAGCACGATGATTTGTTTGCGGGAT<br>GAGCAGTAGCTGCGGGAAACTGCAGATAGTGGTCTGCCC<br>CTGTGGGGGTTGGCAAGTAAGTTGTGGGCCTATCAGCTG<br>GTAGTTAGGGTAATGGCCTAACTAGGCGCAGACGGGATA<br>CGGGGGATCAGGGTTTGGTTCCGGAGAGGGAGCCTGAGA<br>GATGGCTACTACGTCCAAGGATGGCAGCAGGCGCGAAAC<br>TTGCCTAATCCTTTGGGGAGGCGGTTATGAGAAGTGATGT<br>GTGTGCGAGTGCAAAGGGGTCGCATGTGATTGGAGGGCA<br>AGTCGGGTGCCAGCAGCCGCGGTAATACCTGCTCCAATA<br>GTGTCTATGGTGGATGCTGCAGTTAAAATGTCCGTAGTCT<br>GTTGTGTATGTCTTTGTGTGTGATGTTTGTGGTTGTGTGTG<br>GATGTAGTGATGTGTGTGGCAGAGGACGAGGGGCACTGG<br>ATAGTTGGGCGAGAGGTGAAATGCGAAGACCCTGACTGG<br>ACGAGCGGAAGCGAAGGCTGTGCTCTTGGACTAATGTTG<br>CGATGAAGGACGAAGGCTAGAGGATCGAAATCGATTAGAT<br>ACCGTTTTAGTTCTAGCAGTAAACGATGCCGACTGGACGG<br>GACTGTGTGTGTTGTCCATGAGAAATCTTGAGTATGCGGG<br>TTCTGGGGATAGTATGCTCGCAAGAGTGAAACTTGAAGAG<br>ATTGACGGAAGGACACCACAAGGAGTGGAGTGTGCGGCT<br>TAATTTGACTCAACGCGGGGCAACTTACCGGCTCTGAAGG<br>ATGCCTGTGAGTGCATGGCATGAGGCATGCGGCGGTGGT<br>GCATGGCCGTTTTAAATGGATGGCGTGAGCTTTGTCTTAA<br>GTTGCGTAAGATGTGAGACCCTTTGACGGTGTTCTACGGA<br>GCAAGGAGGGGATGGAAGAGAACAGGTCCGTTATGCCCT<br>GAGATGAGGCGGGCTGCACGCGCACTACGATAGATGGCG<br>CTTCTGCCTGCTGTGAGGGATGAAGCTGTGTAAGGGGCTT<br>CTGAACGTGGAATTCCTAGTAATAGCGGCTGACGAAGCTG<br>CTTTGAATGTGTCCCTGTCCTTTGTACACACCGCCCGTCG<br>CTATCTAAGATGACGCACTGGACGAAGATCGGAAGGTCTG<br>AGTCCTGAGTGTTAGATAAGATATAAGTCGTAACATGGCT<br>GCTGTTGGAGAACCAGCAGCAGGATCAGTAT |

**Table S4: NCBI NT accession numbers of additional contextual species included in phylogenetic tree.**

| Accession | Species name |
|---|---|
| JQ062988 | *Ichthyosporidium weissii* |
| HM626203 | *Loma salmonae* |
| GQ203287 | *Glugea hertwigi* |
| AJ252958 | *Pleistophora sp.* |
| MT006314 | *Glugeidae sp.* |
| MW077214 | *Fusasporis stethaprioni* |
| AF356223 | *Ovipleistophora mirandellae* |
| KX099692 | *Pleistophora beebei* |
| GU183263 | *Dasyatispora levantinae* |
| KC137548 | *Heterosporis sp.* |
| AJ002605 | *Trachipleistophora hominis* |
| XR552272 | *Vavraia culicis* |
| AY530532 | *Myosporidium merluccius* |
| KX364284 | *Hyperspora aquatica* |
| KU163282 | *Paradoxium irvingi* |
| DQ417114 | *Thelohania butleri* |

| | |
|---|---|
| HM140491 | *Myospora metanephrops* |
| AY958070 | *Nadelspora canceri* |
| MN935433 | *Ameson herrnkindi* |
| KX856426 | *Perezia nelsoni* |
| HM800849 | *Facilispora margolisi* |
| MF429927 | *Microsporidium sp.* |
| AF356222 | *Kabatana takedai* |
| MH911629 | *Inodosporus octosporus* |
| MF974572 | *Microsporidia sp.* |
| AF364303 | *Tetramicra brevifilum* |
| AY033054 | *Microgemma caulleryi* |
| GQ868443 | *Microsporidia sp.* |
| EU534408 | *Potaspora morhaphis* |
| MG708238 | *Apotaspora heleios* |
| AJ438959 | *Dictyocoela cavimanum* |
| MW377751 | *Unikaryon panopei* |
| JQ268567 | *Triwangia caridinae* |
| GQ206147 | *Neoflabelliforma aurantiae* |
| DQ675604 | *Euplotespora binucleata* |
| GU130406 | *Helmichia lacustris* |

| | |
|---|---|
| MN595900 | *Globosporidium paramecii* |
| FJ914315 | *Mrazekia macrocyclopis* |
| AY233131 | *Cystosporogenes legeri* |
| JX915758 | *Anostracospora rigaudi* |
| L39109 | *Endoreticulatus schubergi* |
| GU130407 | *Crispospora chironomi* |
| AF394525 | *Glugoides intestinalis* |
| GU126383 | *Anisofilariata chironomi* |
| FJ389667 | *Paranucleospora theridion* |
| HG005137 | *Obruspora papernae* |
| U78176 | *Nucleospora salmonis* |
| JX101917 | *Enterospora nucleophila* |
| L07123 | *Enterocytozoon bieneusi* |
| HE584635 | *Hepatospora eriocheir* |
| JX915760 | *Enterocytospora artemiae* |
| KT762153 | *Globulispora mitoportans* |
| KX757849 | *Parahepatospora carcini* |
| KX424959 | *Pancytospora epiphaga* |
| LC136798 | *Percutemincola moriokae* |
| EU709818 | *Liebermannia covasacrae* |

| | |
|---|---|
| KX360142 | *Enteropsectra longa* |
| AJ302316 | *Orthosomella operophterae* |
| FJ865223 | *Mockfordia xanthocaeciliae* |
| KC172651 | *Sporanauta perivermis* |
| AF394529 | *Ordospora colligata* |
| DQ996241 | *Vairimorpha necatrix* |
| KR704648 | *Rugispora istanbulensis* |
| AF495379 | *Oligosporidium occidentalis* |
| MT510137 | *Nosema bombycis* |
| EU275200 | *Heterovesicula cowani* |
| EU075347 | *Binucleata daphniae* |
| KT950767 | *Agglomerata cladocera* |
| DQ641245 | *Senoma globulifera* |
| MK053815 | *Pseudoberwaldia daphniae* |
| AF439320 | *Gurleya daphniae* |
| AF394527 | *Larssonia obtusa* |
| MH645035 | *Conglomerata obtusa* |
| AY090042 | *Berwaldia schaefernai* |
| AY090067 | *Hazardia milleri* |
| AY326268 | *Trichotuzetia guttata* |

| | |
|---|---|
| KX832080 | *Lanatospora costata* |
| AY090041 | *Marssoniella elegans* |
| AY880951 | *Paraepiseptum polycentropi* |
| AY880953 | *Episeptum circumscriptum* |
| KT950766 | *Alfvenia sibirica* |
| KC990122 | *Multilamina teevani* |
| FN794114 | *Octosporea muscaedomesticae* |
| EF537880 | *Zelenkaia sp.* |
| JF826402 | *Amblyospora bakcharia* |
| HM594269 | *Trichotosporea pygopellita* |
| AF027684 | *Edhazardia aedis* |
| AY013359 | *Intrapredatorus barri* |
| AY326269 | *Culicospora magna* |
| AF027683 | *Culicosporella lunata* |
| AF483837 | *Hyalinocysta chapmani* |
| EU664450 | *Andreanna caspii* |
| JF826419 | *Novothelohania ovalae* |
| AY090065 | *Parathelohania obesa* |
| KF110990 | *Takaokaspora nipponicus* |
| AJ252962 | *Flabelliforma montana* |

| | |
|---|---|
| AY090069 | *Polydispyrenia simuli* |
| AF132544 | *Caudospora palustris* |
| KR704917 | *Myrmecomorba nylanderiae* |
| EF564602 | *Ovavesicula popilliae* |
| XR001214623 | *Nematocida parisii* |
| GU173849 | *Kneallhazia carolinensae* |
| AF024658 | *Tubulinosema acridophagus* |
| MF278272 | *Fibrillaspora daphniae* |
| AY364089 | *Fibrillanosema crangonycis* |
| AY953292 | *Systenostrema alba* |
| AY135024 | *Schroedera plumatellae* |
| MN512229 | *Neoperezia semenovaiae* |
| AF484691 | *Bryonosema plumatellae* |
| AF484695 | *Trichonosema pectinatellae* |
| MN752317 | *Jirovecia sinensis* |
| AJ581995 | *Bacillidium vesiculoformis* |
| AF484694 | *Pseudonosema cristatellae* |
| JX463178 | *Pseudonosematidae sp.* |
| AY305324 | *Paranosema locustae* |
| AF024655 | *Antonospora scoticae* |

| | |
|---|---|
| NG017174 | *Rozella allomycis* |

**Text S1: Newick file format of generated phylogenetic tree.**

(SRR17317295_Vairimorpha_ceranae:0.0216381998,(((((((((SRR23560257_Encephalitozoon_hellem:0.1270261617,SRR24007515_Encephalitozoon_intestinalis:0.0579490963)100:0.0456705591,SRR17858635_Encephalitozoon_cuniculi:0.1222391079)100:0.1689587732,(FJ865223_Mockfordia_xanthocaeciliae:0.2351313696,KC172651_Sporanauta_perivermis:0.1712250092)100:0.1447538512)99:0.1045040235,AF394529_Ordospora_colligata:0.3556069511)100:0.1865243865,(((((SRR16954902_Hamiltosporidium_magnivora:0.0012993579,SRR16954899_Hamiltosporidium_tvaerminnensis:0.0011732195)100:0.5406977425,((((((((SRR23214363_Nematocida_ausubeli:0.0482534697,SRR17622377_Nematocida_major:0.0493801792)76:0.0611071277,XR001214623_Nematocida_parisii:0.1849040428)100:0.4652495597,EF564602_Ovavesicula_popilliae:0.4426099381)100:0.3477337868,NG017174_Rozella_allomycis:0.6754674174)78:0.1068215572,(((((ERR3154977_Tubulinosema_ratisbonensis:0.0000022854,AF024658_Tubulinosema_acridophagus:0.0067459583)100:0.1750434162,(SRR489790_Anncaliia_algerae:0.1447156518,GU173849_Kneallhazia_carolinensae:0.0889520487)100:0.2731873561)100:0.0731958285,(MF278272_Fibrillaspora_daphniae:0.1766558011,AY364089_Fibrillanosema_crangonycis:0.0741322584)53:0.0285987202)100:0.2426575762,AY953292_Systenostrema_alba:0.5208555303)100:0.1984565831,(((((AY135024_Schroedera_plumatellae:0.0919222332,MN512229_Neoperezia_semenovaiae:0.0397933062)100:0.0556154110,AF484691_Bryonosema_plumatellae:0.0824629792)100:0.1716363648,((MN752317_Jirovecia_sinensis:0.1158401129,AJ581995_Bacillidium_vesiculoformis:0.0909695828)99:0.0327068725,AF484694_Pseudonosema_cristatellae:0.0802172379)85:0.0184956767)55:0.0155306582,AF484695_Trichonosema_pectinatellae:0.1914992208)89:0.0612273043,(JX463178_Pseudonosematidae_sp.:0.0386666509,(AY305324_Paranosema_locustae:0.0739440307,AF024655_Antonospora_scoticae:0.0674201252)100:0.3546640985)85:0.0951921410)84:0.0737498620)94:0.1198060800)85:0.0540200851,(((AJ252962_Flabelliforma_montana:0.0063553719,AY090069_Polydispyrenia_simuli:0.0000025293)100:0.0704156265,AF132544_Caudospora_palustris:0.0361128181)100:0.2562863668,KR704917_Myrmecomorba_nylanderiae:0.2908173305)97:0.0999217655)62:0.0652994279,((((((((((EU075347_Binucleata_daphniae:0.0154618938,KT950767_Agglomerata_cladocera:0.0017020810)100:0.0198998135,DQ641245_Senoma_globulifera:0.0246503852)100:0.1133531132,(MK053815_Pseudoberwaldia_daphniae:0.0190169136,AF439320_Gurleya_daphniae:0.1134753681)99:0.0166662804)100:0.0403159600,(AF394527_Larssonia_ob tusa:0.0156522832,(MH645035_Conglomerata_obtusa:0.0014183543,AY090042_Berwaldia_schaeferNai:0.0102258848)97:0.0022384546)100:0.0410537983)100:0.0820119073,((AY090067_Hazardia_milleri:0.0264256589,AY326268_Trichotuzetia_guttata:0.6340574444)100:0.0465786137,KX832080_Lanatospora_costata:0.1549169278)80:0.0294242096)81:0.0221325200,((AY090041_Marssoniella_elegans:0.0667540110,AY880951_Paraepiseptum_polycentropi:0.0694903241)86:0.0171228013,AY880953_Episeptum_circumscriptum:0.0408303421)100:0.0530717028)82:0.0240123901,(KT950766_Alfvenia_sibirica:0.1348622419,KC990122_Multilamina_teevani:0.2348299510)100:0.0914857820)93:0.0261180393,(FN794114_Octosporea_muscaedomesticae:0.0310447358,EF537880_Zelenkaia_sp.:0.1517445864)100:0.1957642542)100:0.1008378497,(((((JF826402_Amblyospora_bakcharia:0.0721882307,HM594269_Trichotosporea_pygopellita:0.0876401701)100:0.0708278561,(AY013359_Intrapredatorus_barri:0.1085863613,AY326269_Culicospora_magna:0.2060430602)99:0.0393414688)96:0.0240137948,AF027684_Edhazardia_aedis:0.1043646946)100:0.0685470076,(AF027683_Culicosporella_lunata:0.2422390995,AF483837_Hyalinocysta_chapmani:0.1871064858)100:0.2000473950)100:0.1524032155,EU664450_Andreanna_caspii:0.1312590309)100:0.1321396606)89:0.0370114608,((JF826419_Novothelohania_ovalae:0.2345871757,AY090065_Parathelohania_obesa:0.1025986450)98:0.0475389834,KF110990_Takaokaspora_nipponicus:0.2135945994)100:0.3169546912)100:0.3225655619)93:0.0732389839,SRR8476225_Astathelohania_contejeani:0.9716926742)80:0.0320719677)62:0.0273459869,GQ206147_Neoflabelliforma_aurantiae:0.4225018617)100:0.1147984091,(((((((SRR8495097_Cucumispora_dikerogammari:0.0958231988,KX364284_Hyperspora_aquatica:0.0454922737)100:0.1295947359,KU163282_Paradoxium_irvingi:0.0753711397)100:0.0346847578,DQ417114_Thelohania_butleri:0.0527039767)100:0.0623539816,HM140491_Myospora_metanephrops:0.0967847710)100:0.0836304788,(((((((((SRR926312_Pseudoloma_neurophilia:0.0952068544,JQ062988_Ichthyosporidium_weissii:0.0677985901)100:0.0579158214,HM626203_Loma_salmonae:0.1465261306)100:0.0643191452,GQ203287_Glugea_hertwigi:0.0896394524)96:0.0168636949,(MT006314_Glugeidae_sp.:0.1680443775,MW077214_Fusasporis_stethaprioni:0.0516842256)77:0.0094881930)72:0.0097538182,AJ252958_Pleistophora_sp.:0.1164831586)100:0.0395965512,(((AF356223_Ovipleistophora_mirandellae:0.0346709337,KX099692_Pleistophora_beebei:0.0274897012)100:0.0265606136,(GU183263_Dasyatispora_levantinae:0.0571063667,KC137548_Heterosporis_sp.:0.0551893922)100:0.0231861705)100:0.0469799171,(AJ002605_Trachipleistophora_hominis:0.0394630502,XR552272_Vavraia_culicis:0.0333540041)100:0.0779474640)98:0.0169302239)100:0.0879966826,AY53

0532_Myosporidium_merluccius:0.1407055378)100:0.1002103729,((AJ438959_Dictyocoela_caviman um:0.1027783275,MW377751_Unikaryon_panopei:0.1331711850)100:0.1306302717,JQ268567_Triwangia_caridinae:0.1943026206)71:0.0181541751)59:0.0204207869,((((AF356222_Kabatana_takedai:0.0676219601,MH911629_Inodosporus_octosporus:0.0538168112)99:0.0195582672,MF974572_Microsporidia_sp.:0.1065315768)100:0.0380554533,((AF364303_Tetramicra_brevifilum:0.0131319767,AY033054_Microgemma_caulleryi:0.0267823213)100:0.0564370255,GQ868443_Microsporidia_sp.:0.0426940700)100:0.0530918074)95:0.0255011920,(EU534408_Potaspora_morhaphis:0.0970586148,MG708238_Apotaspora_heleios:0.0619594883)100:0.1314275782)100:0.0926865154)86:0.0640042105)85:0.0544900999,((AY958070_Nadelspora_canceri:0.0096415889,MN935433_Ameson_herrnkindi:0.0295780956)100:0.7853415148,KX856426_Perezia_nelsoni:0.3602093328)88:0.1964533677)73:0.0503818248,(HM800849_Facilispora_margolisi:0.2670456534,MF429927_Microsporidium_sp.:0.2615014740)100:0.2668789765)100:0.3304866548)100:0.1799322487,((((((((SRR23214350_Pancytospora_philotis:0.0800866087,(KX424959_Pancytospora_epiphaga:0.0510661795,LC136798_Percutemincola_moriokae:0.0374253143)100:0.2168517810)100:0.1079356232,AJ302316_Orthosomella_operophterae:0.0837423593)90:0.0774093314,KX360142_Enteropsectra_longa:0.1782780486)97:0.0628781992,EU709818_Liebermannia_covasacrae:0.3511045746)100:0.2703574964,(((((((((SRR065293_Vittaforma_corneae:0.0716005919,(JX915758_Anostracospora_rigaudi:0.1273622596,L39109_Endoreticulatus_schubergi:0.1011501623)100:0.0876087293)52:0.0225323769,AF394525_Glugoides_intestinalis:0.2866656568)83:0.0308668002,GU130407_Crispospora_chironomi:0.0781307271)48:0.0357242504,AY233131_Cystosporogenes_legeri:0.0855683638)94:0.0382588863,FJ914315_Mrazekia_macrocyclopis:0.0901358378)95:0.0339202792,MN595900_Globosporidium_paramecii:0.0100668014)95:0.0594784048,(DQ675604_Euplotespora_binucleata:0.0832730518,GU130406_Helmichia_lacustris:0.3236589665)100:0.0730791450)100:0.3558603737,SRR926341_Agmasoma_penaei:0.5902622021)97:0.0416218281,GU126383_Anisofilariata_chironomi:0.3776814544)88:0.0624766202)94:0.0471809403,(JX915760_Enterocytospora_artemiae:0.0739895299,KT762153_Globulispora_mitoportans:0.0510152278)100:0.1587117348)65:0.0211585274,KX757849_Parahepatospora_carcini:0.1618536433)73:0.0469466533,((((FJ389667_Paranucleospora_theridion:0.1763596231,HG005137_Obruspora_papernae:0.1565482355)96:0.0287010097,U78176_Nucleospora_salmonis:0.0832004481)100:0.0901402672,(JX101917_Enterospora_nucleophila:0.1454874602,L07123_Enterocytozoon_bieneusi:0.2072854861)100:0.0568614394)100:0.0387261408,HE584635_Hepatospora_eriocheir:0.35

08670384)100:0.0772828820)100:0.2920163588)100:0.1773260440)99:0.1802956633,EU275200_Heterovesicula_cowani:0.7346172373)100:0.4783502727,MT510137_Nosema_bombycis:0.2859775967)100:0.1499259805,AF495379_Oligosporidium_occidentalis:0.0251747795)95:0.0192353697,DQ996241_Vairimorpha_necatrix:0.0359240190)100:0.0167039222,KR704648_Rugispora_istanbulensis:0.0412102351);

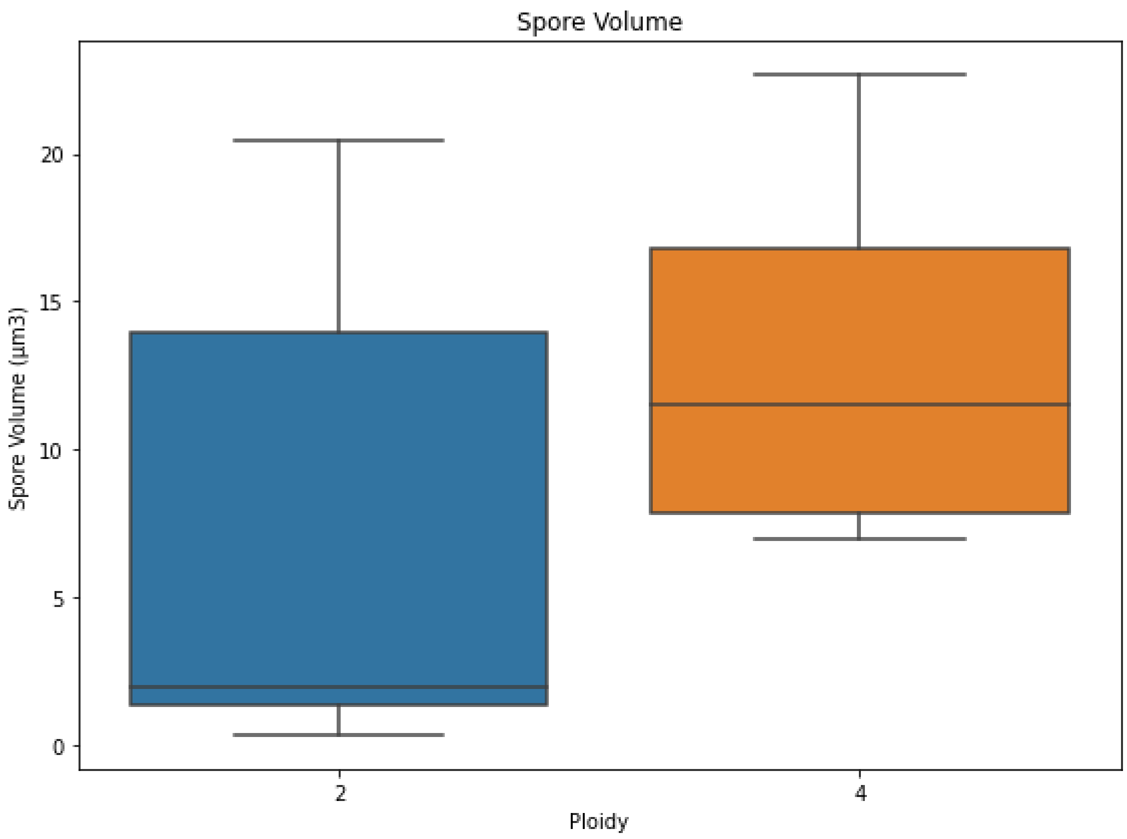


**Figure S1: Box plot of spore volume in diploid and tetraploid species, calculated using data from Bojko et al. (2022).**

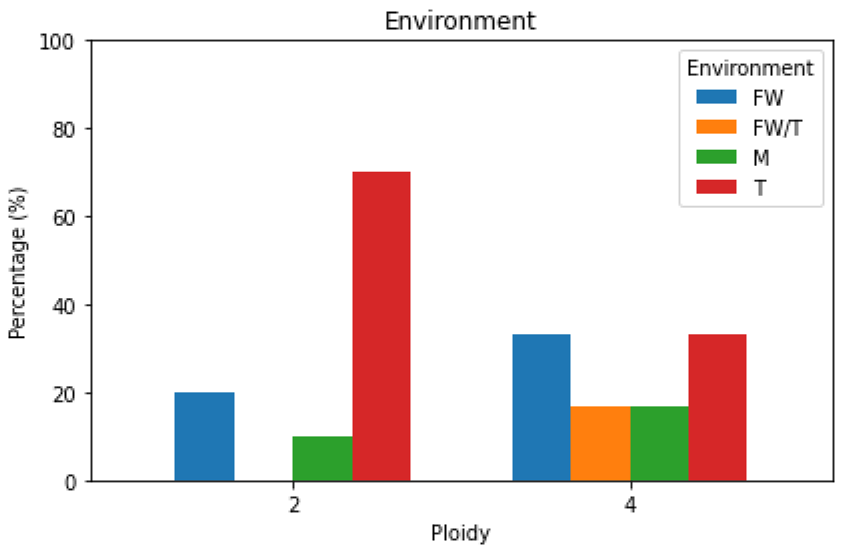


**Figure S2: Bar plot of habitats the diploid and tetraploid species identified in this study are found in according to Bojko et al. (2022).** FW: Fresh water, T: terrestria, and M: marine.

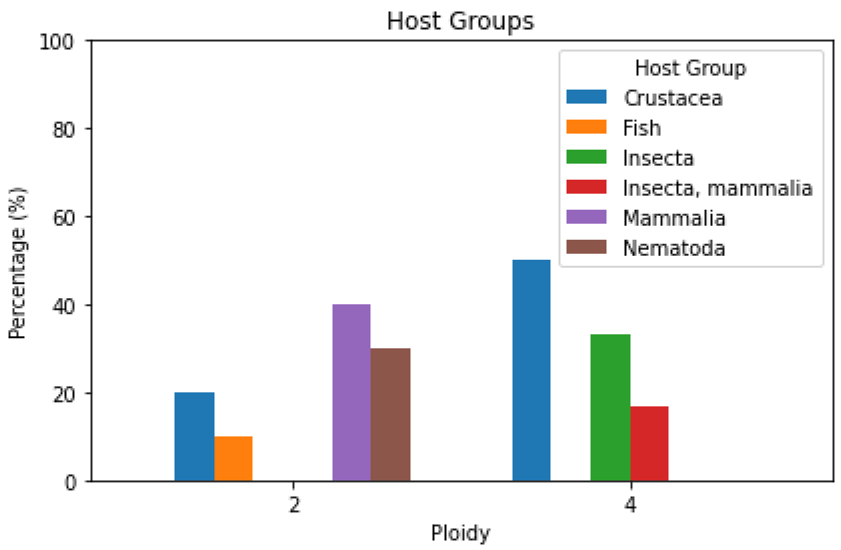


**Figure S3: Bar plot of different host groups the diploid and tetraploid species identified in this study occur in, according to Bojko et al. (2022).**

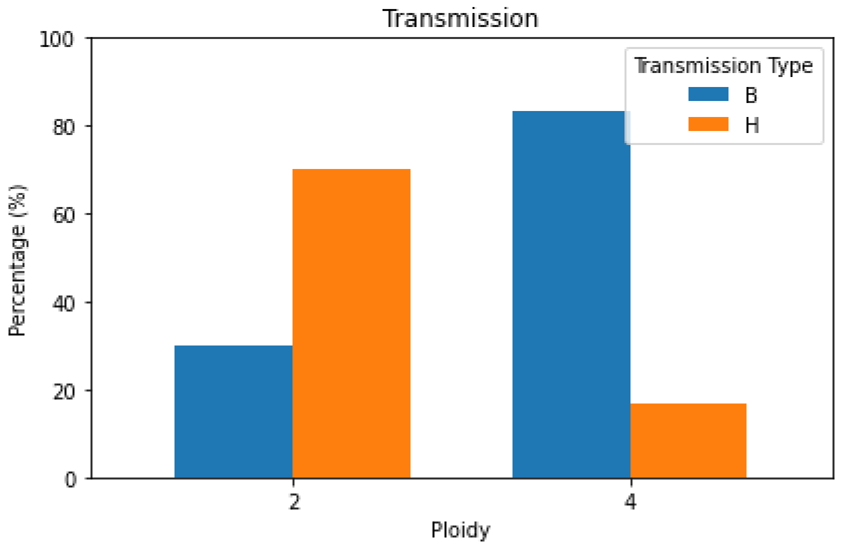


**Figure S4: Bar plot of different transmission modes the diploid and tetraploid species identified in this study possess, according to Bojko et al. (2022).** H: Horizontal, V: Vertical, B: Both.

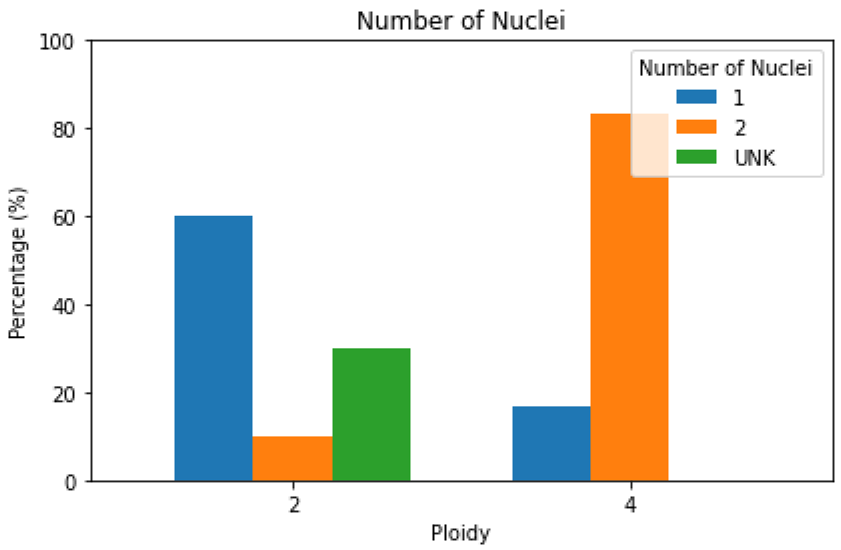


**Figure S5: Bar plot of number of nuclei in the spores of the diploid and tetraploid species identified in this study, according to Bojko et al. (2022).** UNK: Unknown.
